# Mitotic polarity oscillation promotes epithelial tumor progression

**DOI:** 10.1101/2025.02.06.636979

**Authors:** Gayaanan Jeyanathan, Ming Meggie Cao, Milena Pellikka, Sarah Robinson, Parama Talukder, Vanessa Ghorayeb, Ulrich Tepass

**Affiliations:** Department of Cell and Systems Biology, University of Toronto; Toronto, ON M5S 3G5, Canada

**Author notes:** Correspondence should be addressed to: Ulrich Tepass, Department of Cell and Systems Biology University of Toronto, 25 Harbord Street, Toronto, Ontario M5S 2M6 Canada.

## Abstract

Mitosis of epithelial cells requires a transient loss of epithelial polarity [1–5]. However, the nature of this mitotic polarity oscillation and its functional consequences for epithelial development are not fully understood. Here we show that the Crumbs (Crb) complex, a key regulator of epithelial polarity, is lost from the membrane during mitosis, and the Crb mutant phenotype is ameliorated when cell division is inhibited. Remarkably, an essential requirement of Crb for epithelial polarity is fully suspended when cell division is blocked in conjunction with inhibition of either cell ingression or cell intercalation. We conclude that the amount of morphogenetic stress induced by mitosis, ingression, and intercalation determines the requirement for Crb. Increased cell division and loss of cell polarity are two main drivers of epithelial cancer [6–8]. Maintaining epithelial polarity is important for limiting proliferation. Whether the loss of polarity during mitosis impacts tissue growth is less clear. We show that increasing cell division in a morphogenetically quiet epithelium not only increases tissue size but also causes hyperplastic to neoplastic transition. Conversely, reducing cell division restores epithelial polarity in neoplastic tissue of tumor mutants. Taken together, our study revealed that a major function of polarity factors in epithelial maintenance is to counteract morphogenetic stress. Moreover, we propose a feedforward mechanism that links cell division and the loss of polarity as a key driver of epithelial cancer.

---

Mitosis is accompanied by a dramatic cytoskeletal reorganization including the formation of a uniform actin-based cytocortex to facilitate cell rounding, the microtubule-based mitotic spindle to power chromosomal alignment and segregation, and an actomyosin ring that coordinates cytokinesis [2,3,9–11]. In the context of solid tissues such as epithelia, mitosis appears incompatible with maintaining normal tissue architecture. Epithelial integrity depends on apical-basal polarity of individual cells, a polarity that involves an asymmetric cytoskeleton and associated adhesion complexes. Polarity is governed by a network of polarity proteins which include the apical Par and Crumbs (Crb) complexes and basolateral factors such as Par1, Yurt, and Scribbled (Scrib) [12–14]. Indeed, epithelial polarity is transiently lost during mitosis to allow changes in cell shape and cytoskeletal organization required for mitosis, and apical-basal polarity is re-established at the end of mitosis [4,5]. This ‘mitotic polarity oscillation’ has the potential to disrupt epithelial organization. However, direct evidence for the functional consequences of mitosis on epithelial integrity is lacking.

A large majority of cancer cases are derived from epithelial cells. Compromised epithelial polarity is one important aspect of tumor progression [7,8,15]. A second central hallmark of tumor formation is enhanced cell proliferation, raising the possibility that the mitotic polarity oscillation makes important contributions to tumor progression. We assessed the relationship between mitosis and epithelial polarity in the context of normal development and neoplastic tumors. We first asked how factors that regulate apical-basal polarity behave during mitosis and how mitosis affects epithelial integrity. We report that the Crb complex is lost during mitosis and re-expressed during the subsequent interphase, and demonstrate that mitosis challenges epithelial polarity, enhancing the requirement for the activity of the epithelial polarity machinery. Moreover, cell division cooperates with cell ingression and cell intercalation to challenge the epithelial phenotype. We next asked whether the frequency of cell division can impact tumor progression. We discovered that an increase in proliferation rate not only enlarges the tissue but also disrupts epithelial integrity causing the formation of neoplastic tumors. Finally, we asked whether reducing proliferation affects neoplastic tumors. We found that tumors are smaller, as expected, when cell division is reduced but also that epithelial organization is restored despite the loss of key polarity factors such as Scrib. Based on this evidence we propose that neoplastic tumor formation is not initiated by the loss of polarity leading to enhanced proliferation but originates in enhanced proliferation which through the associated mitotic polarity oscillation collapses epithelial structure, fostering tumor progression.

## Transient loss of Crb and Sdt during epithelial mitosis

The transmembrane protein Crb is a component of the apical polarity network and a key regulator of epithelial polarity in the Drosophila embryo [12,16,17]. In a screen for genes that control Crb protein distribution or levels, we found that the Drosophila Cdc20 homolog (*fizzy*; *fzy*) is required to maintain normal Crb levels in the ventral and head epidermis (Fig. 1A,B). Cdc20 acts as an activator of the anaphase promoting complex [18]. *fzy* mutant embryos are depleted of maternally provided gene product at the end of gastrulation at which point cells undergo metaphase arrest, predominantly in the ventral and head ectoderm. Crb surface levels were strongly reduced in metaphase arrested cells of *fzy* mutants (Fig. 1A,B). In addition to Crb, also several other apical markers were reduced or redistributed in metaphase arrested cells, including the Crb binding partner Stardust (Sdt), the apical Par complex proteins atypical Protein Kinase C (aPKC) and Par6, and the apical cadherin Cad87A (Fig. S1). Markers of adherens junctions such as E-cadherin (Ecad) and Par3 (Bazooka [Baz] in Drosophila) appeared somewhat reduced and more fragmented (Fig. S1). In contrast, basolateral markers including Discs Large (Dlg), Yurt, and the Na^+^K^+^-ATPase remained associated with the membrane at relatively normal levels (Figs. 1 A,B; S1). Together, these results suggest that Crb and other apical factors are depleted form the apical surface of epithelial cells undergoing metaphase arrest.

**Figure 1:**
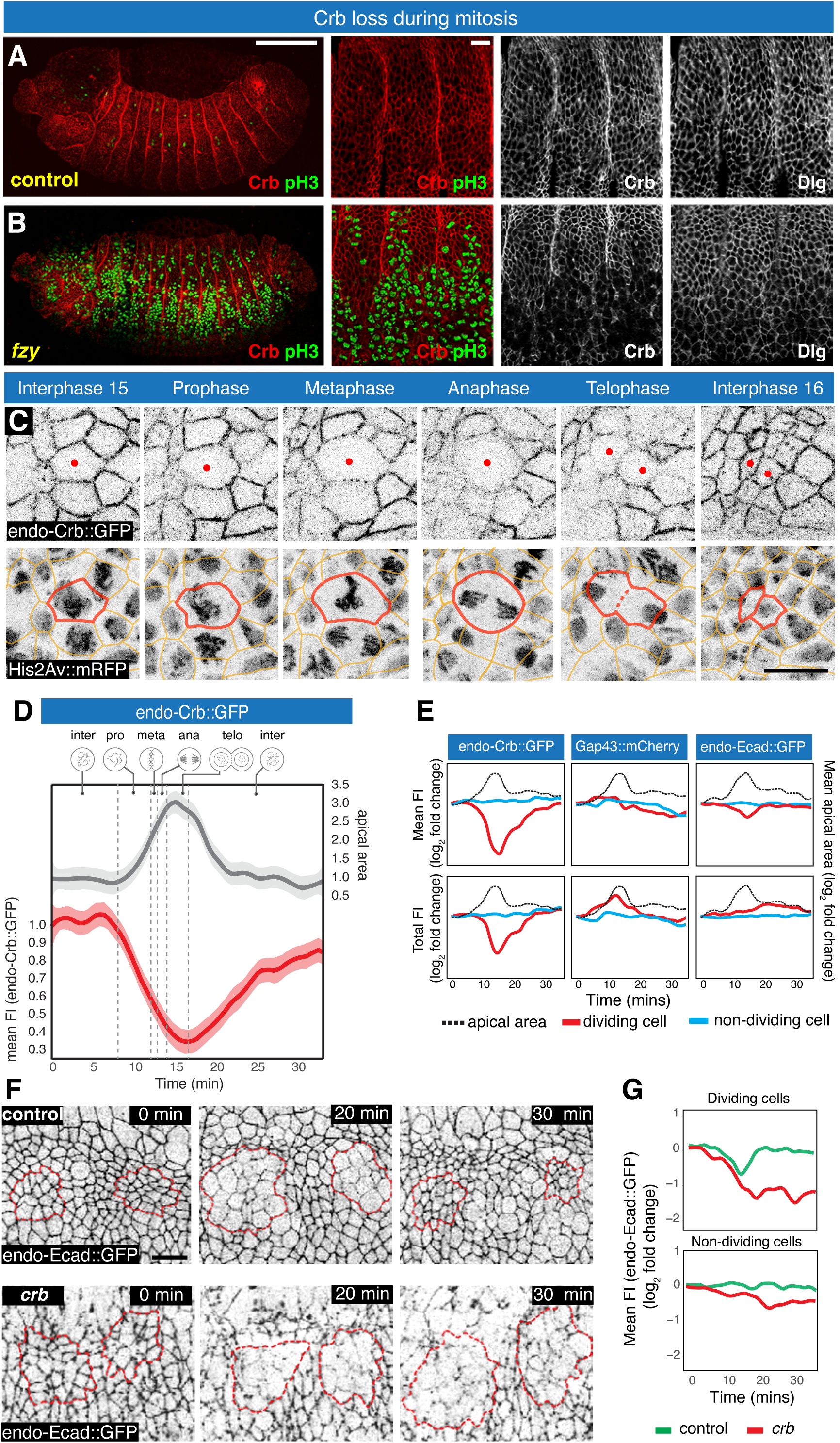
Crb is transiently lost during mitosis. (**A-B**) Wild-type (A) and *fzy* mutant (B) embryos at stage 14 stained for Crb (red), Dlg, and the mitosis marker pH3 (green). Scale bar, 100 μm (left panel) and 10 μm (right panels). Metaphase arrested cells in the *fzy* mutant embryo show reduced Crb levels. (C) Stills of a time-lapse movie showing dividing cells in mitotic domain 8_15_11 of the ectoderm expressing endo-Crb::GFP and the chromosomal marker His2Av::mRFP. Red dots and outline, respectively, highlight a dividing cell. Scale bar, 5 μm. (D) Mean fluorescence intensity (FI) of endo-Crb::GFP (red) and apical cell area (grey) during cell division. Starting values were normalized to 1. (E) Mean FI and total FI of endo-Crb::GFP, Gap43::mCherry, and endo-Ecad::GFP during cell division. (F) Stills of time-lapse movies of endo-Ecad::GFP in control and *crb* mutant embryos undergoing their second postblastodermal cell division in the dorsal ectoderm. Clusters of dividing cells are outlined in red. Scale bar, 10 µm. (G) Mean FI of endo-Ecad::GFP in dividing and non-dividing cells in the 8_15_11 mitotic cluster of control (green) and *crb* mutant (red) embryos. N values are listed in Table S2

To assess the behavior of Crb during mitosis in live wild-type embryos we used an endogenously GFP-tagged Crb protein (endo-Crb::GFP) [19]. Focussing on the second post-blastodermal mitotic division [20], we found that Crb surface levels rapidly declined during prophase while cells underwent mitotic rounding and rose to the apical site of the epithelium (Fig. 1C, D; videos S1 and S2). Crb levels continued to decline during meta- and anaphase with the lowest level seen in telophase. Crb increased during the subsequent interface to reach normal levels ∼6-7 minutes after the completion of telophase. A similar loss of apical distribution was observed for Sdt (Fig. S2; video S3).

We compared protein dynamics of endo-Crb::GFP with GAP43::mCherry, a general membrane marker, and endo-Ecad::GFP, which labels adherens junctions. Interphase cells maintained uniform protein levels for all three markers during the observation period (Fig. 1E). GAP43::mCherry showed a moderate increase during mitotic rounding in conjunction with the enlargement of the apical cell area. This together with the strong decrease of Crb during mitosis indicates that the loss of Crb is not caused by apical area increase during mitotic rounding. In contrast to Crb, total Ecad levels per cell remained constant during mitosis. However, Ecad mean fluorescent intensity decreased during early mitosis suggesting that the increase in perimeter during mitotic rounding lowers junctional Ecad concentration (Figs. 1E and S1). Similar to Crb, Ecad levels recovered after mitosis along the perimeter of the two daughter cells. Our findings support the view that epithelial polarity oscillates during cell division, with a transient loss of apical-basal polarity [4,5].

The cell surface and the cytoskeleton undergo a dramatic reorganization during prophase to prepare an epithelial cell for division. The adherens junctions that link epithelial cells into a sheet are retained during division [5,21–24]. However, the adherens junction-associated actomyosin cytoskeleton disassembles. Actin reorganizes into a uniform cytocortex that supports cell rounding during prophase, whereas myosin II assembles at the equator of the cell during anaphase, where the cytokinetic furrow will form. Epithelial polarity proteins are also temporally reassigned. Proteins of the apical Par complex (Cdc42, aPKC, Par6) and the apical FERM domain protein Moesin redistribute from the apical membrane to the entire cell surface where they promote the formation of a mitosis-specific actin cortex, which is crucial for cell rounding and normal division [4,5,25–28]. As the Crb-complex stabilizes apical aPKC/Par6, Moesin and the epithelial adherens junctions in interphase cells [12,14,29], it is possible that the loss of Crb during mitosis has an important permissive role in facilitating cytoskeletal reorganization. The re-appearance of Crb at the apical membrane at the end of mitosis suggests that Crb is involved in the re-polarization of postmitotic epithelial cells. Using adherens junction integrity (endo-Ecad::GFP) as a readout for normal polarity, we found that adherens junctions reassemble was severely compromised after division in *crb* mutants (Fig. 1F, G). These findings suggest that Crb is required to establish apical-basal polarity in daughter cells as they reform normal epithelial polarity at the end of mitosis.

## Mitosis disrupts epithelial integrity

Embryos that lack Crb lose epithelial integrity during gastrulation when epithelial cells undergo three rounds of cell division, including the loss of apical markers, fragmentation of adherens junctions, and loss of the sheet-like epithelial tissue structure [16,20,30,31]. Peak disruption of epithelial integrity is observed at the end of gastrulation (embryonic stage 11). In post-gastrulation embryos, many epithelial cells in *crb* mutants activate JNK signaling and undergo programmed cell death. The remaining epithelial cells, however, re-polarize and organize into small epithelial cysts with the apical side facing inward [29,31,32,33]. Indeed, when cell death is blocked all epithelial cells in *crb* mutants form epithelial cysts suggesting that Crb is dispensable for epithelial polarity in post-gastrulation embryos [31]. Thus, an essential requirement for Crb in maintaining epithelial polarity overlaps with the time period when epithelial cells undergo cell division.

To test whether Crb is required to support epithelial polarity in the absence of cell division we examined *crb* compromised embryos where cell division was blocked during gastrulation using mutations of *string* (*stg*), which encodes the Drosophila cell cycle regulator Cdc25 [34]. Blocking cell division partially rescued the epithelial defects cause by the loss of Crb. (Fig. 2D,F), suggesting that Crb function in epithelial polarity is more important when cells proliferate.

**Figure 2:**
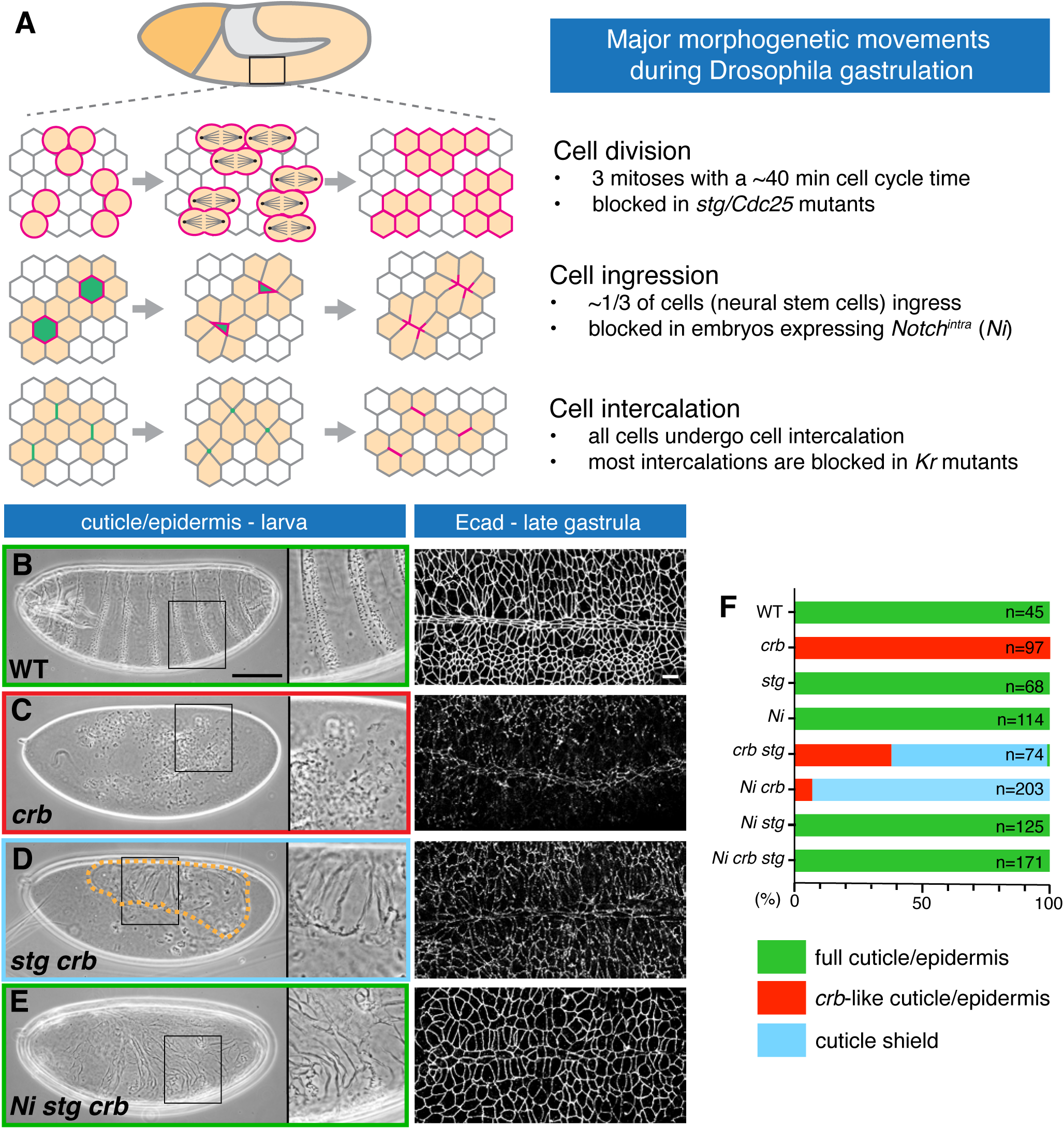
Reducing morphogenetic stress ameliorates epithelial defects in *crb* mutant embryos. (A) The trunk ectoderm undergoes three main morphogenetic movements during gastrulation: cell division, cell ingression and cell intercalation which can be selectively blocked by the indicated genetic tools. (**B-E**) Larval cuticle and embryo at late gastrulation (stage 11) labelled with endo-Ecad::GFP in control (WT), *crb*, *stg crb*, and *Ni stg crb* mutants. Scale bars, 100 µm (cuticle) and 10 µm (Ecad staining). Orange dotted line highlights a cuticle shield in a *stg crb* mutant embryo. We distinguished between animals that are fully surrounded by cuticle/epidermis (green frame; B,E), animals showing a cuticle shield (blue frame; D), and embryos with a *crb*-like cuticle fragmentation (red frame; C). (**F**) Quantification of cuticle/epidermal defects in larvae of the listed genotypes. N values are shown in Table S2.

A partial rescue of epithelial integrity was observed previously in embryos compromised for Ecad or Cdc42 in which epithelial cell ingression was reduced [35,36]. We also observed a partial rescue of epithelial defects in *crb* mutants when cell ingression was blocked (Fig. 2F; S3E). Like cell division, the ingression of neural stem cells (neuroblasts) during gastrulation requires the formation of new cell contacts that need to be polarized to maintain epithelial integrity (Fig. 2A) [37]. We therefore hypothesized that cell division and ingression pose a similar challenge to epithelial polarity. To test this, we blocked in *crb* mutants both cell division (by loss of Stg/Cdc25) and the ingression of neuroblasts through the expression of Notch^intra^ (Ni), an active form of the Notch receptor that inhibits neuroblast specification [38] (*Ni stg crb* embryos). Remarkably, the *crb* mutant phenotype was fully suppressed in *Ni stg crb* embryos (Figs. 2E,F and S3). Adherens junctions, which were strongly disrupted in *crb* mutants, were partially restored in *stg crb* embryos and fully restored in *Ni stg crb* embryos at the end of gastrulation (Fig. 2B-E). Similarly, the epidermis/cuticle of *Ni stg crb* embryos was indistinguishable from *Ni stg* embryos (Figs. 2E,F and S3). While these embryos showed many patterning defects because of compromised N and Stg functions, they have an intact epidermal epithelium that fully surrounds the embryo as in wild-type. In contrast, *Ni crb* embryos only showed a moderate amelioration of the *crb* phenotype similar to *stg crb* embryos (Figs. 2D,F and S3). These data suggest that both cell division and cell ingression challenge epithelial integrity and that these processes operate additively in enhancing ‘morphogenetic stress’. Crb loses its essential function in epithelial polarity when morphogenetic stress is substantially reduced, emphasizing the context-dependent nature of Crb function in polarity maintenance.

A third morphogenetic process that is observed during gastrulation in addition to cell division and cell ingression is the extension of the germband, which requires extensive cell intercalation (Fig. 2A) [39]. To reduce cell intercalation in a Crb compromised embryo we took advantage of mutations in the segmentation gene *Krüppel* (*Kr*) [39]. Interestingly, the loss of *Kr* ameliorates somewhat the *crb* mutant phenotype similar as seen in Crb compromised embryos in which either cell division (*stg crb* embryos) or cell ingression (*Ni crb* embryos) was blocked [40]. Moreover, inhibiting either cell division in *Kr crb* embryos (*Kr stg crb* embryos) or cell ingression (*Ni Kr crb* embryos) caused a striking rescue of the *crb* polarity phenotype (Figs. S3H,I and S4). Whereas epithelial development appeared fully restored in *crb* mutant embryos with compromised cell division and cell intercalation, moderate defects in the ventral ectoderm persist in *crb* embryos in which cell ingression and cell intercalation was blocked, suggesting that cell division may challenge the epithelium more so than cell ingression (Fig. S4D,E). Epithelial defects in *crb* mutants were suppressed when two of the three major morphogenetic processes observed during gastrulation were blocked, implying that the epithelium can cope with some level of morphogenetic stress in the absence of Crb. Taken together, these findings suggest that cell division, cell ingression, and cell intercalation are functionally similar in challenging epithelial polarity. Consequently, the function of Crb may either be essential or dispensable depending on the level of morphogenetic activity experienced by an epithelium.

Crb cooperates with the apical Par complex and adherens junction proteins such as Ecad in supporting epithelial polarity [12,14], raising the question whether the sensitivity of gene function to cell division is a Crb-specific feature, or a more general aspect of the cell polarity machinery. We tested whether the loss of cell division could rescue epithelial defects in embryos with a compromised Par complex or defective adherens junctions. Cdc42 acts as an apical polarity protein to localize and stimulate the aPKC/Par6 complex during Drosophila gastrulation [36,41]. Compromised Cdc42 (through the expression of a dominant-negative construct, Cdc42-DN) causes defects in the ventral epidermis while the dorsal epidermis remains largely intact [36]. These defects were partially suppressed by blocking cell division in embryos expressing Cdc42- DN (*Cdc42-DN stg* embryos) (Fig. 3A-C and I). As epithelial integrity of the epidermis is only partially disrupted in Cdc42-DN embryos we asked whether an increase in cell division would further disrupt epithelial polarity. Ectopic expression of Stg causes additional cell divisions [34].

**Figure 3:**
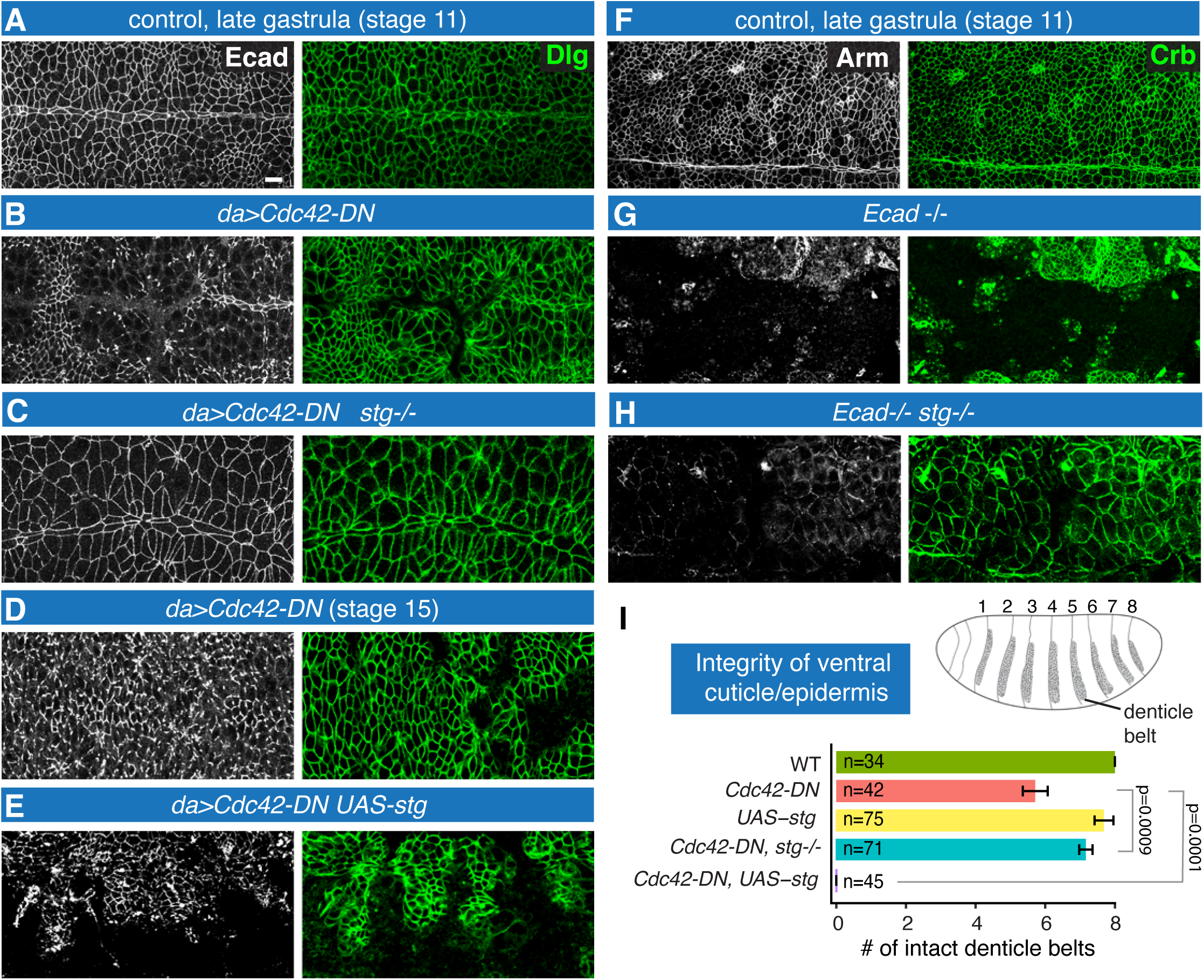
Cell division-induced morphogenetic stress modulates epithelial defects in the *Cdc42* and *Ecad* mutant embryos. (**A-C**) Ectoderm of embryo at late gastrulation (stage 11) labelled for the adherens junction marker Ecad and the basolateral marker Dlg of control (A), *da-Gal4 UAS-Cdc42-DN* (B) and *da-Gal4 UAS-Cdc42-DN stg* (C) genotypes. *da-Gal4* induces ubiquitous expression of target genes. Note restoration of junctional integrity in the *Cdc42-DN stg* embryo that lacks cell division compared to *a Cdc42-DN* embryo. Scale bar, 10 µm. (**D,E**) Epidermis of stage 15 embryos labelled for Ecad and Dlg of the genotypes *da-Gal4 UAS- Cdc42-DN* (D) and *da-Gal4 UAS-Cdc42-DN UAS-stg* (E). Increased cell division enhances epithelial defects. (**F-H**) Ectoderm embryo at late gastrulation (stage 11) labelled for the adherens junction marker Arm and the apical marker Crb of control (F), *Ecad* mutants (G), and *Ecad stg* mutants (H). Zygotic *Ecad* mutant shows a reduction of the Ecad-binding partner Arm (G, H). *Ecad stg* mutants which lack division show a continuous ventral ectoderm compared to the large gaps seen in *Ecad* mutants. (**I**) Quantification of ventral cuticle defects. We counted the number of intact abdominal denticle belts (1-8, see schematic) in embryos of the listed genotypes. Whereas suppression of cell division (*stg-/-*) ameliorates the *Cdc42-DN* phenotype, increased cell division (*UAS-stg*) strongly enhances it. N values are shown in Table S2.

Ectopic expression of Stg in wild type did not cause a loss of epidermal tissue. In contrast, ectopic Stg expression strongly enhanced epidermal defects in Cdc42-DN embryos (Figs. 3D,E,I and S3J,K). This suggests that the requirement of the apical Par complex is sensitive to the amount of cell proliferation. Similarly, epithelial defects that are caused by compromising the adherens junction components Ecad [35] or Baz/Par3 [27] are ameliorated by reducing cell division or a combination of reducing cell division and cell ingression (Figs 3F-I and S3L,M). We conclude that the requirement for the apical epithelial polarity machinery in general is closely linked to the amount of epithelial cell division.

## Mitosis promotes tumor progression

Our findings show that cell division exerts morphogenetic stress that challenges epithelial integrity. Enhanced proliferation and the loss of epithelial polarity are hallmarks of tumor development [6,8]. We asked whether excessive proliferation could be a driver not just for increasing tumor size but also of tumor progression from a hyperplastic (adenoma-like) to a neoplastic (adenocarcinoma-like) state. To address this question, we examined the relationship between cell division and epithelial integrity in larval wing imaginal discs, an established model for the analysis of tissue growth and tumorigenesis [42,43]. One striking difference between the embryo and the larval wing imaginal disc is morphogenetic activity. Whereas the gastrulating embryo experiences a high level of cell division with a rapid cell cycle time of ∼40 minutes [20] as well as cell ingression and cell intercalation in a 2-hour time window, imaginal discs have long cell cycle times of 10-12 hours [44] and undergo comparatively little if any cell ingression or cell intercalation. Interestingly, in contrast to embryos where Crb is essential for epithelial integrity, loss of Crb from imaginal disc epithelia does not compromise epithelial polarity (Fig. 4A-C) [45,46].

**Figure 4:**
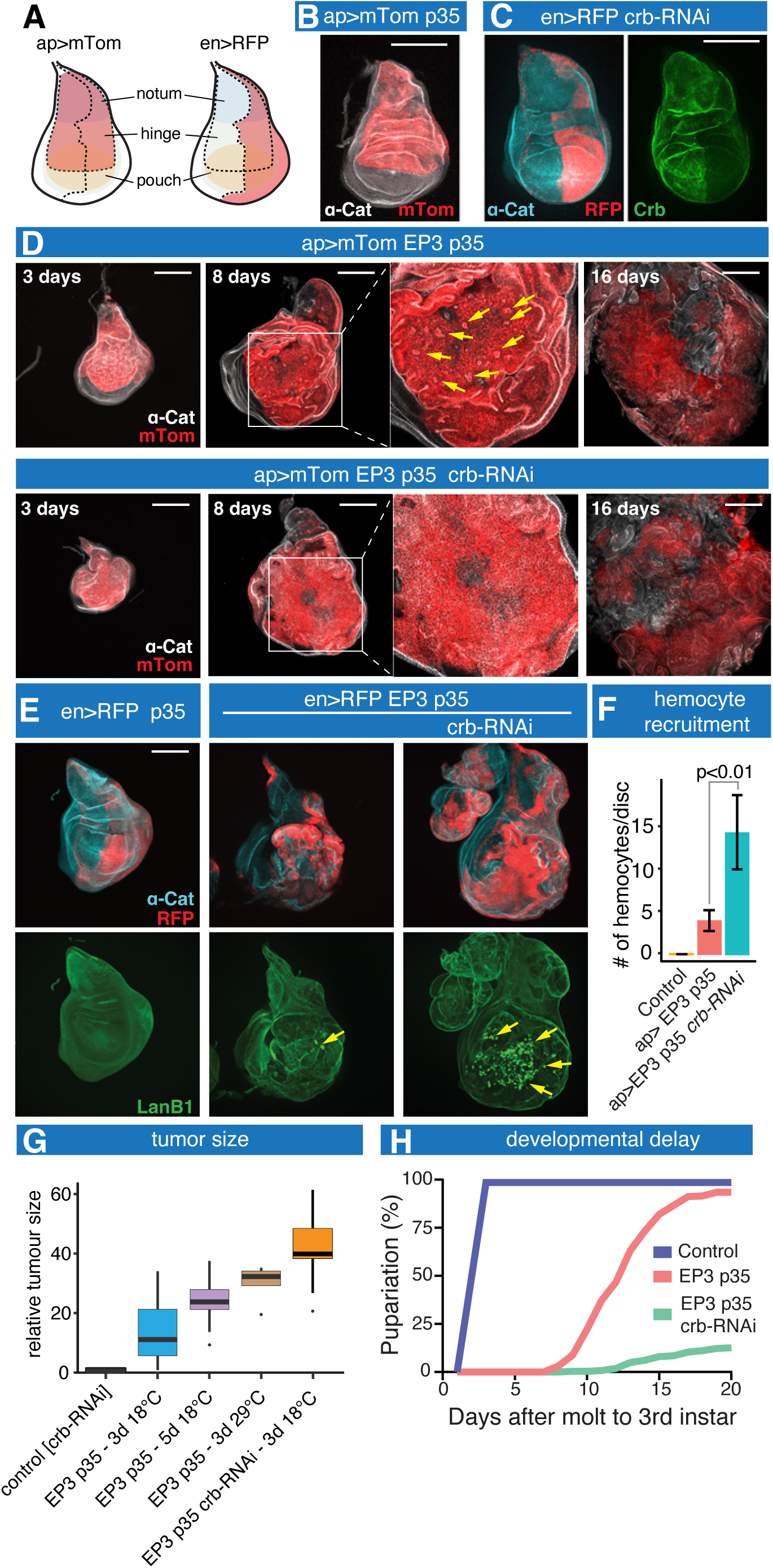
Acceleration of the cell cycle causes neoplastic tumor formation. (A) Expression domains (red) in the wing imaginal disc of *apterous* (*ap*; dorsal compartment) and *engrailed* (*en*; posterior compartment) Gal4 drivers. (B) Wing disc expressing *ap-Gal4 UAS-myrTomato* (*ap>mTom*) *UAS-p35*. Disc is labeled for ⍺- Cat. (C) Wing disc expressing *en-Gal4 UAS-RFP* (*en>RFP*) *UAS-crb-RNAi*. Disc is labeled for ⍺-Cat and Crb. Note depletion of Crb in the posterior compartment. (D) Discs expressing *ap>mTom EP3*(=*UAS-E2F1^PIP-3A^*) and *UAS-p35* or *ap>mTom, EP3, p35*, and *crb-RNAi* stained for ⍺-Cat. Discs were dissected 3, 8, and 16 days at 18°C after molt to 3 larval instar. Arrows in close-up point to epithelial cysts embedded in a neoplastic matrix. Note enhanced neoplastic progression at 8 days as evident from the lack of epithelial cysts and reduced epithelial folding with *crb-*RNAi expression. (E) *en>RFP p35, en>RFP EP3 p35*, and *en>RFP EP3 p35 crb-RNAi* discs stained for ⍺-Cat and Laminin B1 (LanB1). Discs were dissected 5 days at 29°C after molt to 3 larval instar. Note that Crb knockdown causes an increase in disc size and accumulation of hemocytes (yellow arrow) indicating enhanced neoplastic development. (F) Quantification of hemocyte accumulation is wing discs at 5 days at 29°C after molt to 3 larval instar of the indicated genotypes. (G) Relative neoplastic tumor sizes in wing discs of the indicated genotypes grown at 18°C or 29°C for 3 or 5 days after molt to 3 larval instar. Quantification of relative tumor size is illustrated in Figure S8. (H) Developmental delay in pupariation time (days after molt to 3rd larval instar at 18°C) of *ap>mTom EP3 p35* and *ap>mTom EP3 p35 crb-RNAi* animals compared to control. N values are listed in Table S2. Scale bars, 100 µm.

We first asked whether accelerating cell division could not only increase larval wing discs size but also push epithelia into neoplastic growth. We took advantage of a constitutively active form of the cell cycle regulator E2F1 (E2F1^PIP-3A^; ‘EP3’ in the following) to drive cell cycle progression [47]. E2F1 is a transcription factor that promotes entry into S-phase through upregulation of Cyclin E and other cell cycle factors. EP3 was co-expressed with p35 [48] to block programmed cell death that may occur as a result of enhanced proliferation. 3 days after molt to 3rd larval instar at 18°C, wing discs showed a moderate overgrowth of EP3 p35 expressing cells. Overgrowth was strongly enhanced at 8 days and even more so at 16 days after molt (Fig. 4D). Features of neoplastic growth became apparent as tumor size increased including the loss of epithelial organization, the disruption of the basement membrane and the accumulation of F-actin and Mmp1, the recruitment of hemocytes, and elevated JNK, ROS and Wingless signaling in tumor cells (Figs. 4D-F and S5). [e.g. 42,49-51]. Consistent with the presence of neoplastic tumors we observed a delay in pupariation (Fig. 4H) as seen in other neoplastic tumor mutants [42,52]. Neoplasms elicited by EP3 p35 expression could be further enhanced by raising animals at higher temperature (29°C), taking advantage of the temperature sensitive nature of the Gal4/UAS-driven EP3 expression (Fig. 4G), or by co-expression of additional cell cycle regulators such as Stg or a combination of CyclinD and Cdk4 (Fig. S6). These findings suggest that an increase in proliferation causes neoplastic development consistent with the view that cell division compromises epithelial polarity.

Notably, knockdown of Crb in EP3 p35 expressing wing discs accelerated overgrowth and neoplastic development (Fig. 4D). The difference was apparent at 3 and 8 days post molt whereas at 16 days post molt EP3 p35 and EP3 p35 *crb*-RNAi animals showed large neoplasms of comparable size (Fig. 4D,G). The growth dynamics of EP3 p35 *crb*-RNAi wing discs is similar to discs in mutants of classical neoplastic tumor suppressor genes. Wing discs of these mutants are initially smaller than normal 3 days post molt to 3^rd^ larval instar, when wild-type larvae undergo pupariation but then grow dramatically during an extended larval period [42,53]. Enhanced neoplastic tumor growth of EP3 p35 *crb*-RNAi compared to EP3 p35 discs was also evident from the increased recruitment of hemocytes, a signature of neoplastic development (Fig. 4E,F) [51]. Finally, we found that pupariation times were further significantly delayed with EP3 p35 *crb*- RNAi co-expression (Fig. 4H). Together, these data indicate that the loss of Crb enhances neoplastic development in a context of high levels of proliferation. Crb has been linked to the regulation of Notch, JNK, and Hippo signalling and *crb* mutant wing discs show moderate overgrowth while maintaining epithelial polarity [54,55]. The pronounced acceleration of neoplastic development seen in EP3 p35 *crb*-RNAi tissue therefore suggests that Crb function becomes essential for supporting epithelial polarity when cell division frequency is increased.

To further assess the relationship between cell division and tumor progression we reduced proliferation in neoplastic conditions. Neoplastic tumours such as those observed in *scrib* mutant animals over-proliferate and display a loss of epithelial polarity [42,56,57]. Reducing proliferation in tissue that will form a neoplastic tumor is expected to reduce tumor size. Moreover, given the relationship between cell division and epithelial polarity that we discovered, we speculated that reducing proliferation would also restore epithelial integrity despite the loss of Scrib. A moderate reduction of the cell cycle regulator Cdk1 [58] in Scrib-compromised tissue reduced tumor size and, strikingly, restored epithelial polarity (Fig. 5). Crb which displayed a diffuse apolar distribution in Scrib knockdown induced neoplastic tumors showed normal apical localization in *scrib*-RNAi *Cdk1*-RNAi tissue (Fig. 5 C,D). We also examined the distribution of Dlg, a basolateral epithelial polarity factor that cooperates with Scrib and requires Scrib for membrane association [56]. Dlg did not localize to the plasma membrane in Scrib knockdown neoplastic tissue, as expected, but also failed to associate with the membrane in *scrib*-RNAi epithelia that were restored as a result to reduced proliferation (Fig. 5D). This suggests that in *scrib*-RNAi *Cdk1*- RNAi epithelia neither Scrib nor Dlg is required for epithelial polarity. These data support the hypothesis that the loss of polarity is not the primary defect in Scrib-compromised imaginal discs. Instead, these findings argue that excess proliferation elicited by the loss of Scrib is responsible for the collapse of epithelial integrity and, thus, neoplastic tumor progression.

**Figure 5:**
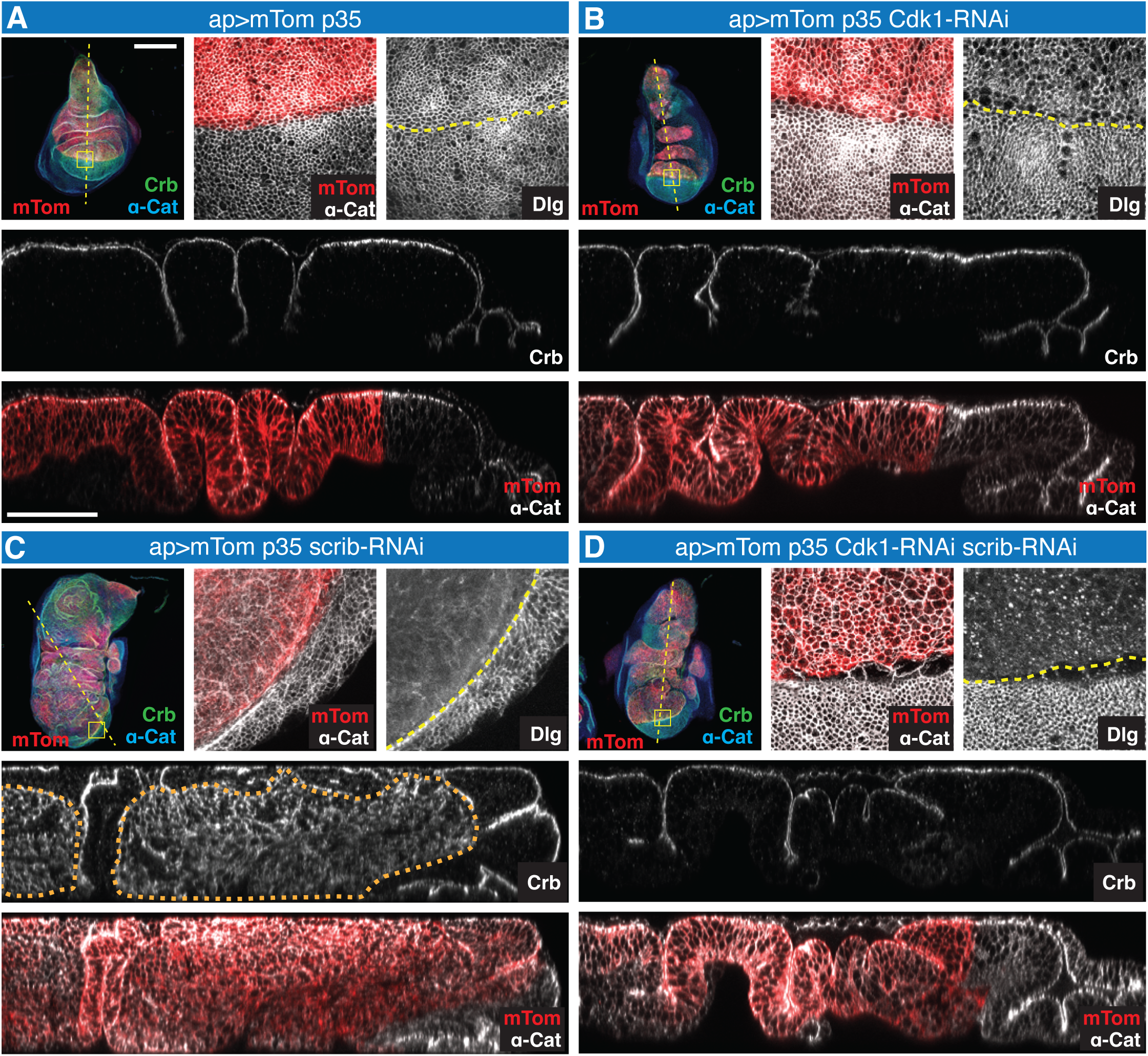
Reducing cell division restores epithelial polarity in neoplastic tumors. (**A-D**) Wing discs expressing *ap-Gal4 UAS-myrTomato* (*mTom*) *UAS-p35* (A), *UAS*-*Cdk1-RNAi UAS-p35* (B), *UAS*-*scrib-RNAi UAS-p35* (C), or *UAS*-*scrib-RNAi UAS-p35 UAS*-*Cdk1-RNAi* (D). All animals also carried *tub-Gal80^ts^*. Animals were grown at 18°C and transferred to 29°C immediately after 2^nd^ to 3^rd^ instar molt to inactivate Gal80^ts^ and dissected 72 hours later. Discs were stained for Crb, ⍺-Cat, and Dlg. Dashed yellow line in disc overview images indicates location of optical cross-sections shown in lower panels. Yellow box shows the region magnified in upper right panels. Yellow dotted line shown in the Dlg labeled tissue indicates the boundary between *ap* expressing dorsal cells and *ap* negative ventral cells. Orange dashed line marks *scrib-RNAi*-induced neoplastic tumor. Dlg is depleted and Crb and ⍺-Cat are mislocalized in *scrib-RNAi* expressing cells (C). α-Cat and Crb expression are restored when cell division is reduced in *scrib-RNAi* tissue, however, Dlg is still depleted (D). Cross-section view show normal Crb apical localization in controls (A,B) and *scrib-RNAi Cdk1-RNAi* discs (D) in contrast to Scrib-depleted tissue (C). Scale bars, 100 μm (disc overview panels); 50 μm (cross-section images). N values are listed in Table S2.

One of the drivers of neoplastic growth in imaginal discs mutant for *scrib* is JNK signalling. JNK normally promotes cell death to, for example, remove damaged cells. However, if cell death is prevented then JNK can foster proliferation [59]. The loss of neoplastic tumor suppressors such as Scrib elicits a strong activation of the JNK pathway [60,61]. The main downstream effector of JNK is the Jak/Stat signaling pathway which promotes cell cycle progression [59,62]. To ask whether JNK is also activated in EP3 p35 expressing tissue we monitored the JNK reporter TRE-dsRed [63]. TRE-dsRed is not detected in wild-type wing discs but strongly expressed in neoplastic EP3 p35 tissue (Fig. S7). In comparison, co-expression of EP3 p35 with *Cdk1*-RNAi caused a reduction in tumor size and TRE-dsRed expression, and a restoration of epithelial organization (Fig. S7). These results further emphasize that the acceleration of the cell cycle causes neoplastic tumor progression similar to the neoplasm observed in mutants of classical neoplastic tumour suppressor genes.

## Discussion

Epithelial polarity is transiently lost during mitosis. Our results show that this mitotic polarity oscillation represents a severe challenge to epithelial integrity. The machinery that maintains epithelial polarity is a complex network of factors the requirement of which can vary from tissue to tissue. We found that the requirement for the polarity machinery depends on the level of morphogenetic stress that is exerted by cell division or two other cell movements, cell ingression and cell intercalation. All three cell behaviours involve changes in cell contacts and the polarization of new cell-cell interfaces [37,64,65], or whole-cell repolarization of the two daughter cells after cell division [4,5, this work]. All three cell movements also exert mechanical stress that can lead to tissue ruptures observed next to dividing or extruding cells in epithelia with weakened cell adhesion or enhanced extrusion [66]. The context dependency - as defined by the degree of morphogenetic activity - of polarity protein requirement explains the striking differences in the strength of epithelial tissue disruption observed in animals carrying mutations affecting polarity proteins such as Crb. Crb is essential for polarity in gastrulating embryos but not required (or redundant) in the late embryonic epidermis or imaginal disc epithelia. Similar findings for the Par complex component Cdc42 or Ecad suggest that this context dependency extends to the entire epithelial polarity machinery and therefore is a fundamental feature of epithelial organization.

Because our data suggested that cell division is a cause of morphogenetic stress in epithelia, we explored the relationship between division and neoplastic growth. Our findings suggest that the mitosis-induced disruption of epithelial polarity is a major driver of tumor progression. Accelerating proliferation in conjunction with blocking cell death in the wing disc epithelium caused neoplastic growth. Conversely, reduction of proliferation rescued neoplastic tumor development and restored epithelial organization of neoplastic tumors. Together, these findings suggest that it is not the loss of epithelial polarity caused by mutations in neoplastic tumor suppressors such as Scrib that is the initiating cause of neoplasm. Instead, our findings argue that cell division and the polarity oscillation during mitosis initiate neoplastic growth as they strongly promote the progression of tumors from a hyperplastic to a neoplastic state. We therefore propose that the primary defect in mutants for *scrib* is an acceleration of proliferation, the detrimental consequences of which may be further enhanced through defects in polarity precipitated by the loss of Scrib and related factors.

Our data argue for a simple model of epithelial tumor progression that entails a feedforward loop between mitosis and epithelial polarity. Components of the polarity machinery are known to limit proliferation. For example, the Crb complex interacts with the Hippo, Notch, and JNK pathways to control proliferation [54,55]. Similarly, epithelial adherens junctions regulate proliferation through several pathways including Hippo and JNK [61,67,68]. Our evidence suggests that high levels of mitosis disrupt epithelial polarity to a degree that severely challenges tissue integrity, a loss of which in turn is predicted to deregulate pathways that limit proliferation. This feedforward mechanisms would be responsive to any signal that enhances proliferation and would then disrupt epithelial polarity which further enhances proliferation, causing persistent neoplastic tissue growth.

## Acknowledgements

We would like to thank Rudi Winkelbauer, Tony Harris, Rodrigo Fernandez-Gonzalez, and Dorothea Godt for critically reading the manuscript and helpful suggestions. We thank Victoria Yan for assistance. We thank many members of the Drosophila research community, the Bloomington Drosophila Stock Center, and the Developmental Studies Hybridoma Bank (University of Iowa) for providing fly stocks and other reagents. We appreciate the support of the Imaging Facility of the Department of Cell and Systems Biology.

## Author contributions

UT designed experiments, analyzed and interpreted data, wrote the manuscript, and raised funding for this project. GJ designed experiments, analyzed and interpreted data and generated the initial draft for this paper. MMC, MP, SR, PT, VG carried out experiments, analyzed and interpreted data. All authors edited the manuscript.

## Data and material availability

All data are available in the main text or the supplementary materials.

## Supplementary materials

Materials and Methods Figures S1 to S8 Tables S1 and S2 Videos S1 to S3

References associated with supplementary materials.

