## Supplemental Methods Figures and Tables for "Mitotic polarity oscillation promotes epithelial tumor progression"

### Supplementary materials

#### Materials and Methods

##### Analysis of Drosophila embryos

**Drosophila genetics:** Flies were raised on standard media supplemented with tetracycline (0.25mg/ml). Fly lines used are listed in Table S1. For genetic interaction experiments that used Gal4 driven overexpression, flies were crossed at 25°C for 24 hours and subsequently moved to 29°C and eggs were collected after 24 hours at 29°C. A full list of genotypes for individual experiments and N values are found in Table S2.

**Preparation of embryonic cuticle**: Embryos were aged for 36–48 hours after egg collection at 25°C, washed, and dechorionated in a 2% bleach solution for 5 min. After washing with double-distilled $H_2O$, eggs were transferred onto a slide into a 1:1 mixture of Hoyer's medium and lactic acid, covered with a coverslip, and incubated overnight at 85°C [69]. Images were taken with a Carl Zeiss Axiophot2 microscope using a phase-contrast 20x lens (NA 0.5). Pictures were recorded with a Canon Rebel XSi camera using Canon software and processed in Adobe Photoshop and Illustrator.

**Embryo fixation and antibody staining:** Drosophila embryos were fixed either with heat or formaldehyde. Embryos were heat fixed by briefly submerging them in 3 ml boiling E-wash (1x E-wash: 4 g NaCl and 0.3 ml Triton X-100 in 1 l $H_2O$) in a 20 ml vial with. ~15 ml of ice-cold E-wash was immediately added. After removing most of the E-wash, embryos were devitellinized by vigorous shaking in a 1:2 heptane:methanol mixture for 30 seconds. For fixation with formaldehyde, embryos were fixed for 20 minutes in 3.7% formaldehyde in a 1:1 PBS:heptane mixture. Embryos were devitellinized as described above. Antibody staining followed standard procedures. Primary and secondary antibodies used are listed in Table S1.

**Imaging and signal intensity quantification:** Leica TCS SP8 scanning confocal microscopes were used to acquire images of live or fixed embryos with 20x and 40x objectives (HC PL APO CS2 with NAs of 1.30 and 1.40, respectively). Time-lapse acquisition protocols were described previously (Simoes et al., 2017). Three to five live embryos were examined for each genotype. Z-stacks were collected using a step size of 0.35-0.45 µm. Stills and videos were assembled from maximum-intensity projections of 12-15 apical planes (ImageJ; National Institutes of Health).

Adobe Photoshop and Adobe Illustrator were used to process and arrange images. The same settings were applied to all images within an experimental series. The average fluorescence intensity of His2AvRFP, endo-Crb::GFP as well as endo-Crb::GFP, GAP43::mCherry and endo-Ecad::GFP, GAP43::mCherry were quantified in segmented cells using Matlab and the script SIESTA (scientific image segmentation and analysis; [70]) and Python3 and the script PyJAMAS [71].

**Statistics**

Statistical analysis was performed in Microsoft Excel or Prism v8 (GraphPad). Data are presented as mean + SEM or mean + SD. Statistical significance was determined using Student's t-test. Experiments were carried out in biological duplicates or triplicates.

**Analysis of Drosophila wing discs**

**Drosophila genetics:** Flies were raised on standard media at 18, 25 or 29°C. A list of fly stocks used is found in Table S1. A *UAS-lacZ* element was added to many control genotypes to maintain a similar number of UAS elements between control and experimental animals. Genotypes for individual experiments and N values are listed in Table S2.

**Developmental series:** To raise larvae at different stages of development, embryos were collected on yeasted 2-inch apple juice agar plates for four hours at 25°C before transfer to 18°C. Upon hatching, larvae were sorted for UAS-myrTomato expression. Larvae were transferred to fresh plates with yeast and maintained at 18°C for 3.5 days. To control for overcrowding and access to nutrients, each plate had a population of 50 larvae. After 3.5 days of incubation, $2^{nd}$ -instar larvae were selected and transferred to fresh plates. These larvae were monitored for 3 hours at which point recently molted $3^{rd}$- instar larvae were selected that were negative for the TM6B balancer (non-Tubby). Molted larvae were identified by morphological changes in the anterior spiracles. The selected batch of $3^{rd}$-instar larvae was then divided into two cohorts, one maintained at 18°C and the second at 29°C. Larvae from both cohorts were dissected every 24 hours which continued for 16 days at 18°C or 12 days at 29°C. 3-5 plates for each genotype were also scored for pupariation every 24 hours.

**Tub-Gal80$^{ts}$ experiments:** Flies were kept at 25°C for 4-hour egg collections on yeasted 2-inch grape juice agar plates after which eggs were incubated at 18°C until hatching. $2^{nd}$-instar larvae that showed a TRE-DsRED signal (an RFP+ signal that is not dependent on Gal4 expression or Gal80$^{ts}$ repression) were sorted and transferred to fresh yeasted plates. Larvae carrying ap-Gal4, UAS-myrTomato were not sorted at this stage due to repression of Gal4 by Gal80$^{ts}$ but were also transferred to fresh yeasted plates. Both cohorts were incubated for 3.5 days at 18°C with a population of 50 larvae per plate. Late $2^{nd}$-instar larvae that were RFP+ and negative for the TM6B balancer (non-Tubby) were transferred to fresh plates and monitored for 1 hour. Freshly molted $3^{rd}$-instar larvae were transferred to 29°C to inactivate Gal80$^{ts}$ allowing the expression of Gal4-driven UAS constructs. Control genotypes were dissected between 36-48 hours after the molt to $3^{rd}$ instar (more than 90% of the larvae had pupariated at 36 hours at 29°C). Genotypes showing disc overgrowth had a pupariation delay and were dissected between 72-96 hours after molt to $3^{rd}$ instar.

**Genetic interaction experiments:** Embryos were collected and maintained at 25°C for 24 hours to allow hatching of $1^{st}$ instar larvae. These larvae were then sorted for RFP+ signal and transferred to fresh yeasted 2-inch grape juice agar plates. Subsequently, $2^{nd}$ instar larvae that were negative for the TM6B balancer (non-Tubby) were selected and transferred to freshly yeasted plates at 50 larvae per plate and monitored every 3 hours for new molted $3^{rd}$ instar larvae. Newly molted larvae were transferred to fresh plates with yeast and once ~70% of the larval population within a plate pupariated, the wing discs of the remaining larvae were dissected and analyzed.

**Immunostaining:** Larval wing imaginal discs were dissected in cold PBS and fixed in 4% paraformaldehyde for 30 min. Discs were washed in PBS containing 0.1% Triton (PBT) 3 times for 10 min and then in 0.3% PBT for 30 min. The tissue was incubated with 10% normal goat serum (NGS) in 0.1% PBT for 2 hours at room temperature before antibody staining. Antibodies used are listed in Table S1. Larval tissues were incubated in the primary antibodies in 10% NGS at 4°C overnight, followed by 4 times 10 min washes in 0.1% PBT. The secondary antibodies were added to larval tissue at room temperature for 2 hours, and DAPI was added to the samples for 15 minutes. The tissue was washed 4 times for 10 min in 0.1% PBT and then 10 min in PBS. Wing discs were mounted either with the apical or basal side up in Vectashield antifade mounting medium (Vectorlabs). A single layer of Scotch double-sided tape was used as spacers to avoid compression of the wing discs.

**Detection of ROS signaling:** Larval wing imaginal discs were dissected in PBS, then treated with CM-H2DCFDA (a general oxidative stress indicator; Table S1) for 15 min in darkness at room temperature. Wing discs were rinsed 3 times with PBS for 5 min and mounted in PBS for live imaging.

**Imaging, image processing and analysis**

Leica TCS SP8 scanning confocal microscopes were used to acquire images of fixed imaginal discs with 20x and 40x objectives (HC PL APO CS2 with NAs of 1.30 and 1.40, respectively). Overview images of imaginal discs are maximum-intensity projections of the top 25 μm of the disc proper (ImageJ; National Institutes of Health). Cross-section views used for quantifications of tumor size were acquired as XZY scans at central locations within the discs using a 40x objective with 0.75x zoom setting. Calculation of relative tumor size is illustrated in Figure S8. Relative tumor size and pupariation data were analyzed using Microsoft Excel and graphed using PrismV8 (Graphpad). Adobe Photoshop and Adobe Illustrator were used to process and arrange images. The same settings were applied to all images within an experimental series.

### Supplemental Figure legends

**Figure S1: Distribution of polarity markers in metaphase arrested cells of *fzy* mutant embryos.**

Epidermis of stage 14/15 *fzy*$^{1}$ mutant embryos with metaphase arrested cells (pH3 marker, green). Apical markers (Sdt, Cad87A, aPKC, and Par6) are strongly depleted in metaphase cells. Adherens junction markers (Ecad and Baz) show a reduction and fragmentation whereas basolateral markers (Dlg, Na/K-ATPase [NaKA] and Yurt [Yrt]) appear normal. Scale bar, 10 µm. N values are listed in Table S2.

**Figure S2: Sdt is transiently lost during mitosis**.

Three stills from video S3 showing the behaviour of endo-Sdt::GFP during the second postblastodermal division of a highlighted cell cluster in mitotic domain 11 [20]. Cells are shown at timepoints 0, 12.5, and 22 min of the video sequence. Note the dramatic loss of Sdt at 12.5 min. N value is listed in Table S2.

**Figure S3: Terminal phenotypes of polarity mutants with reduced morphogenetic stress.**

**(A, A')** Phase contrast image (A) and schematic diagram (A') of cuticle of a wild-type embryo. Orange arrowheads highlight 8 abdominal denticle belts.

(**B-G**) Cuticles of *crb* (B), *Ni* (C), *stg* (D), *Ni crb* (E), *Ni stg* (F) and *Ni stg crb* (G) embryos. Cuticle is largely absent in the *crb* mutant which retains only small specks of cuticle. *Ni* and *stg* mutants have full cuticle. *Ni* mutants fail in germband retraction (arrow points to posterior tip of germband). Cuticle in *stg* mutants is poorly differentiated and faint. Dashed line in (E) surrounds a cuticle sheet indicating a partial rescue of the *crb* mutant phenotype. Note that *Ni stg* and *Ni stg crb* embryos are fully surrounded by cuticle and show similar phenotypes.

(**H,I**) *Kr* (H) and *Kr stg crb* (I) embryos. The *Kr* mutant has a reduced number of denticle belts (orange arrowheads). The *Kr stg crb* embryo is fully surrounded by cuticle. Denticle belts are not differentiated similar to *stg* mutants.

(**J,K)** Embryo expressing Cdc42-DN (J) shows small ventral cuticle defect (dashed line). Embryo expressing Cdc42-DN and overexpressing Stg (K) lacks ventral cuticle (dashed line).

(**L,M**) Embryo expressing BazSA::GFP (L), a mutant form of Baz that cannot be phosphorylated by aPKC, displays a *crb*-like phenotype. BazSA::GFP expressing embryo mutant for *stg* and expressing *Ni* (M) shows full cuticle.
Scale bar, 50 µm. N values are listed in Table S2.

**Figure S4: Loss of cell intercalation and cell division rescues epithelial defects in *crb* mutant embryos.**
(**A-C**) Epidermis of a wild-type (A), *Kr* mutant (B) and *crb* mutant embryo (C) stained for Crb and Ecad.
(**D**) Epidermis of *Ni Kr crb* embryo stained for Crb and Ecad. Orange bracket indicates ventral epidermis which is not fully restored in this mutant.
(**E**) Epidermis of *Kr stg crb* embryo stained for Crb and Ecad. Cells are larger due to the lack of cell division.
Scale bar, 10 µm. N values are listed in Table S2.

**Figure S5: Cell cycle acceleration elicits neoplastic tumor development.**
(**A-E**) Imaginal wing discs expressing *ap-Gal4 UAS-myrTomato* (*mTom*) *UAS-p35* (control) or *ap-Gal4 UAS-mTom UAS-EP3 UAS-p35*. Discs are labeled for Wg to monitor Wg signaling (A), H2-DCF to monitor ROS signaling (B), and Mmp1 to monitor JNK signaling (C). Discs were stained with Phalloidin to detect F-actin enrichment (D), and for Laminin B1 (LanB1) and Hemes to detect basement membranes and hemocytes (E). Note gaps in the basement membrane (yellow arrows) and hemocyte accumulation (red arrowheads). d, days after $2^{nd}$ to $3^{rd}$ larval instar molt; scale bars, 100 µm. N values are listed in Table S2.

**Figure S6: EP3 inducted neoplasm is enhanced by co-expression of other cell cycle regulators.**
(**A-E**) Third instar wing imaginal discs with *ap-Gal4* driving the expression of *UAS-myrTomato* (*mTom*) together with the *UAS-p35* (A), *UAS-p35 UAS-stg* (B), *UAS-EP3 UAS p35* (C), *UAS-EP3 UAS-p35 UAS-stg* (D), *UAS-EP3 UAS-p35 UAS-CycD UAS-Cdk4* (E). Discs were stained for α-Cat to mark apical adherens junctions and DAPI to highlight nuclei. Blue traces show apical α-Cat staining in the mTom positive regions of the discs to indicate the extent of normal epithelial differentiation. Larvae were grown at 29$^{o}$C and dissected when >50% of larvae had pupariated.

*UAS-lacZ* was added to EP3 p35 flies (A and C) to maintain similar numbers of UAS constructs in control and experimental genotypes. Scale bars, 100 µm (in disc overview), 50 µm (in cross-section view). N values are listed in Table S2.

**Figure S7: Reducing cell division lowers JNK signaling and restores epithelial integrity in EP3 p35 wing imaginal discs.**

(**A-D**) *ap-Gal4* driving *TRE-dsRed UAS-p35* (A), *TRE-dsRed UAS-p35 UAS-Cdk1-RNAi* (B), *TRE-dsRed UAS-EP3 UAS-p35* (C), and *TRE-dsRed UAS-p35 UAS-EP3 UAS-Cdk1-RNAi*. Animals also expressed *Tub-Gal80$^{ts}$*, and larvae were grown at 18$^{o}$C until 2$^{nd}$ to 3$^{rd}$ instar molt and then transferred to 29$^{o}$C to inactivate Gal80$^{ts}$. Control discs were dissected at 48 hours post molt (A,B). EP3 p35 expressing discs with and without *Cdk1-RNAi* were dissected at 96 hours post molt (C,D). Depletion of the apical marker Cad87A (cyan; indicating a loss of epithelial polarity) and expression of TRE-dsRed (red, indicating the upregulation of JNK signaling) seen in EP3 p35 tissue (C) is suppressed by co-expression of *Cdk1-RNAi* which reduces proliferation (D). *UAS-lacZ* was added (A and C) to maintain similar numbers of UAS constructs in control and experimental genotypes. Scale bar, 100 µm. N values are listed in Table S2.

**Figure S8: Measurement of relative tumor size.**

**(A)** Schematic of wing disc showing *ap-Gal4* driven *UAS-myrTomato* (*mTom*) expression (red) in a top view (left) and cross-section view (right). Main regions of the wing disc are the pouch, hinge, and notum. *ap* expression is confined to the dorsal compartment.

**(B)** XZY optical cross-section of a wing disc (position indicated by a green dashed line in A) labelled for mTom and α-Cat containing an EP3 p35 induced tumor. To determine relative tumor size, we identified the tumor in the pouch region (yellow outline; arrow points to boundary between pouch and hinge) and measured its area. We then measured (i) the entire apical length of the tumor within the identified tumor region (area facing the lumen between peripodial membrane and disc proper; yellow line in B') and (ii) the length of intact epithelium as identified by normal α-Cat staining (yellow line in B''). Relative tumor size was determined by multiplying the apical length of the tumor (apical length of tumor=yellow line in B'minus yellow line in B'') with the tumor area measured in B.

**Video S1:** Wild-type embryo expressing endo-Crb::GFP and His2Av::mRFP.

**Video S2:** Wild-type embryo expressing endo-Crb::GFP (same embryo as in vídeo S1).

**Video S3:** Wild-type embryo expressing endo-Sdt::GFP.

**Additional references associated with supplementary materials.**

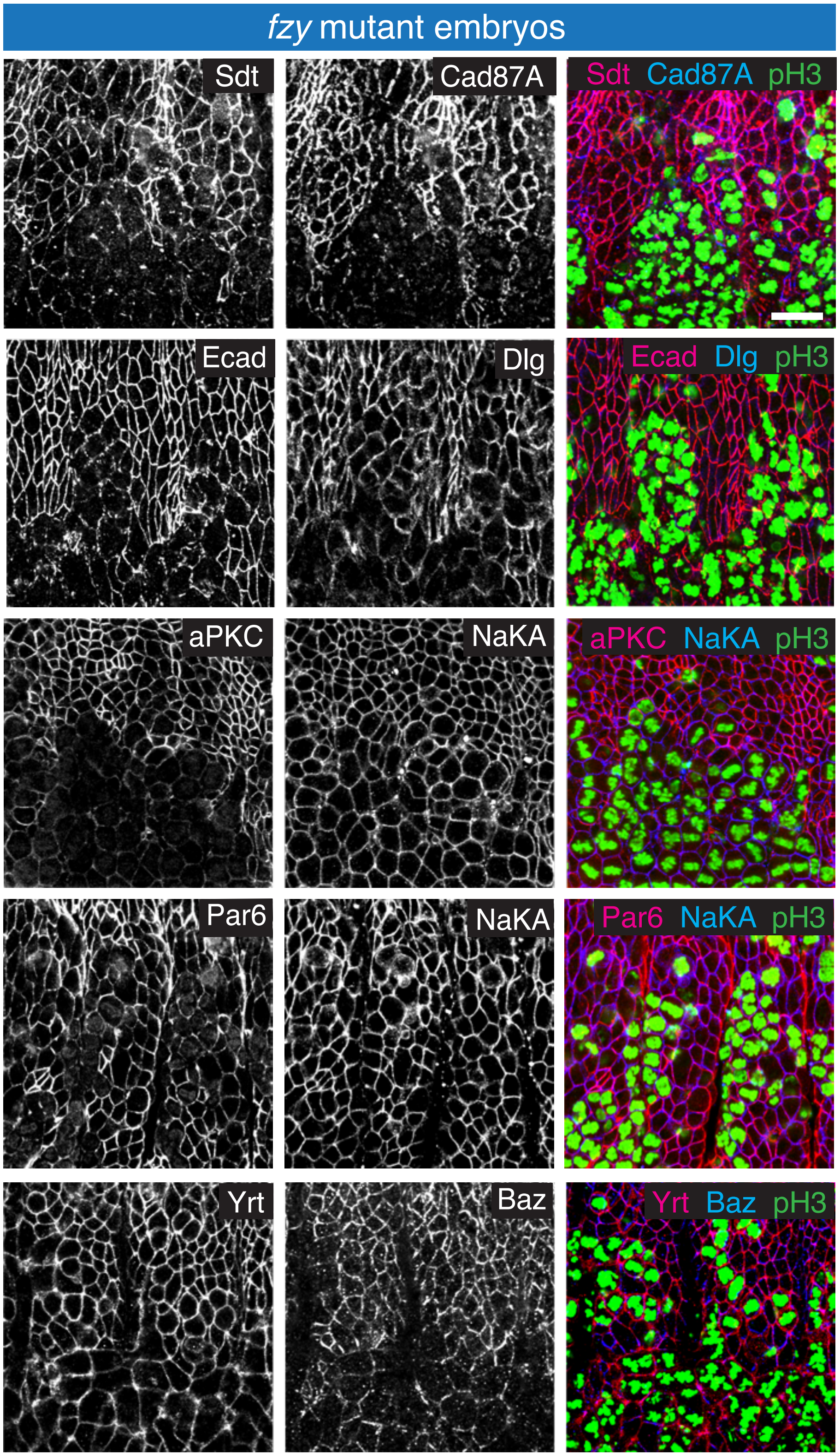

**Jeyanathan et al. Figure S1**

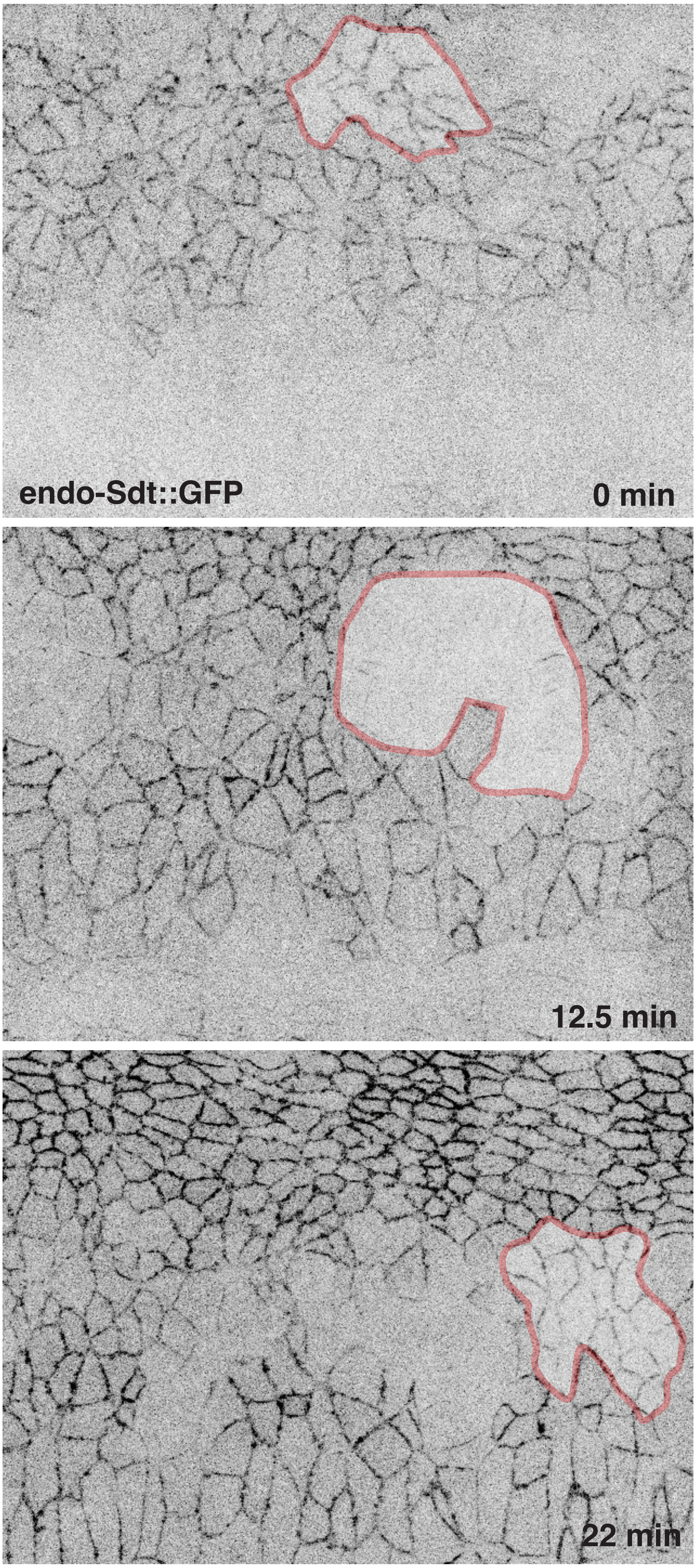

Jeyanathan et al. Figure S2

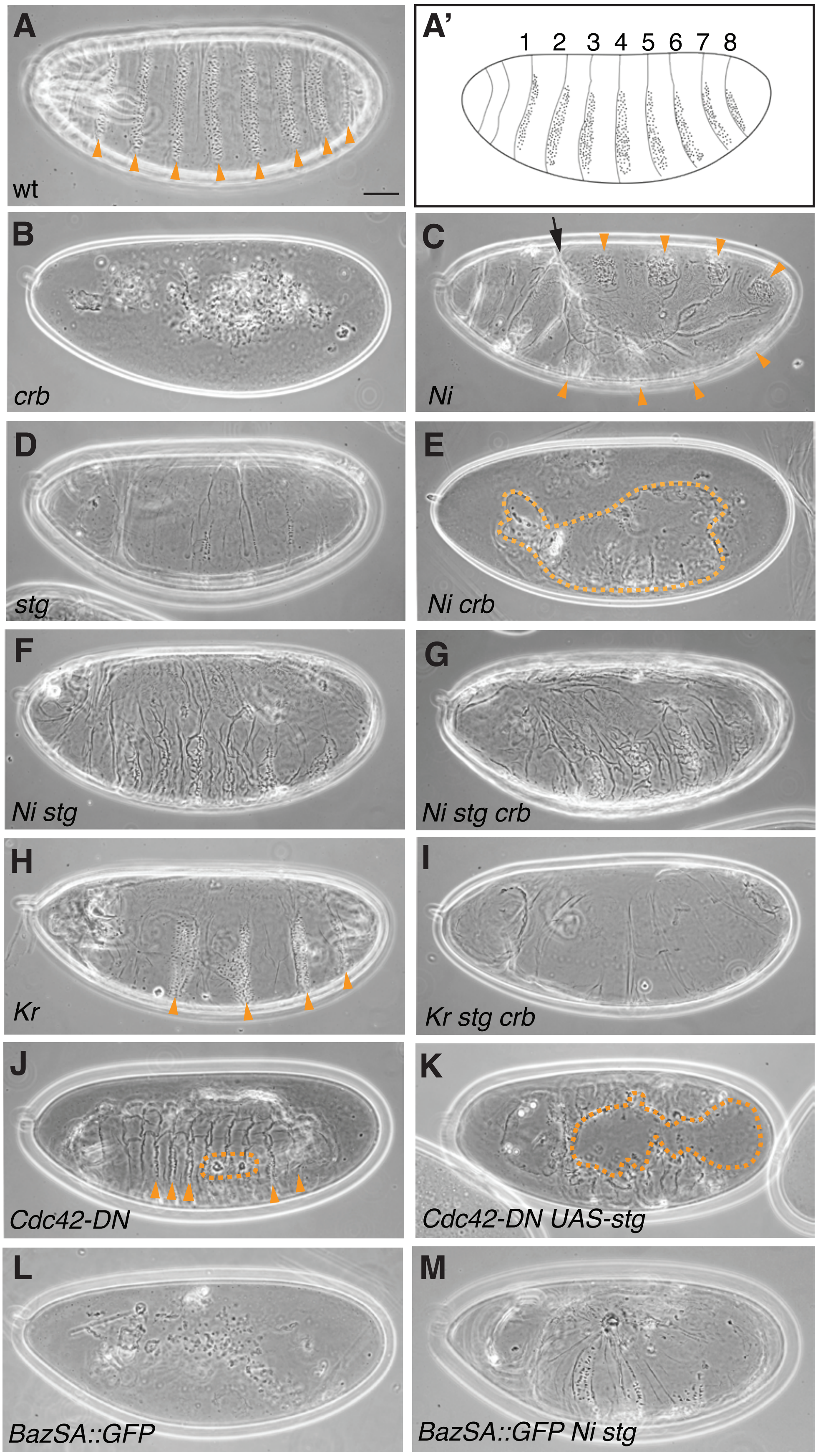

**Jeyanathan et al. Figure S3**

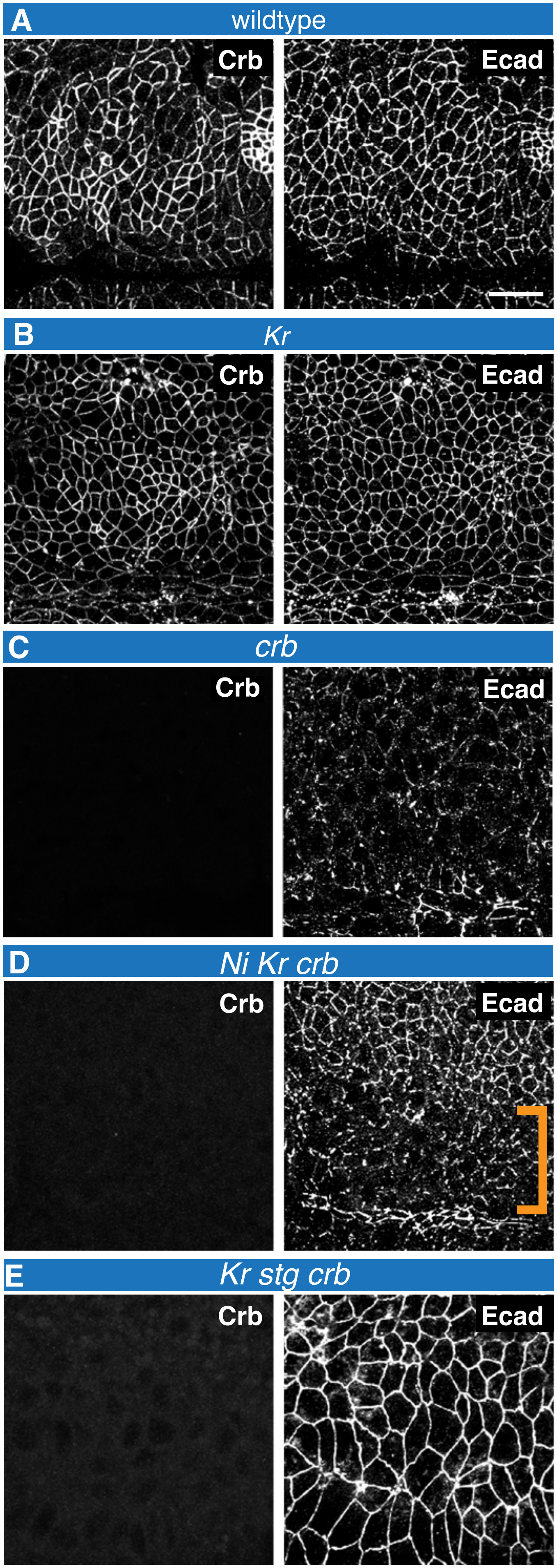

**Jeyanathan et al.** Figure S4

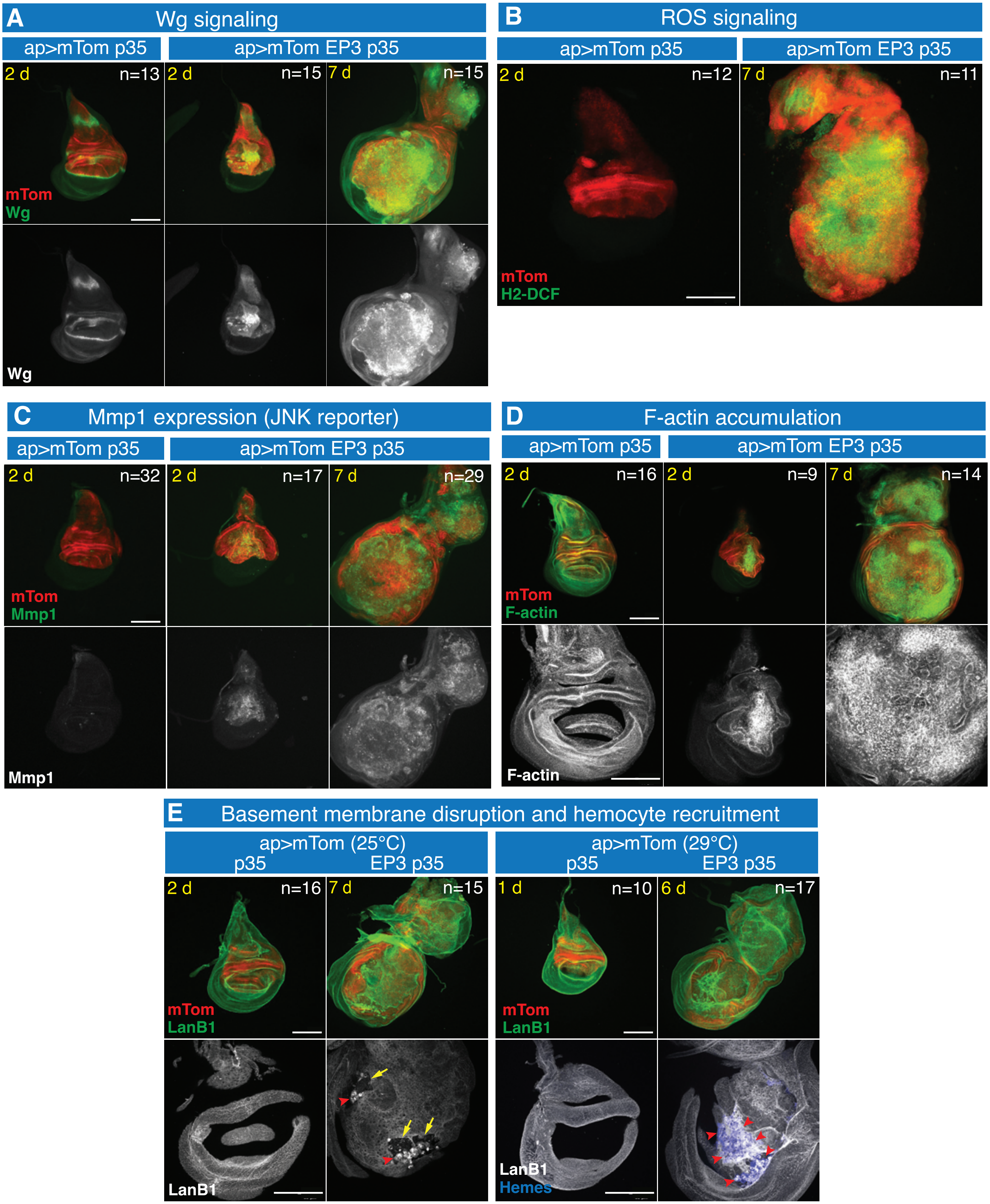

Jeyanathan et al., Figure S5

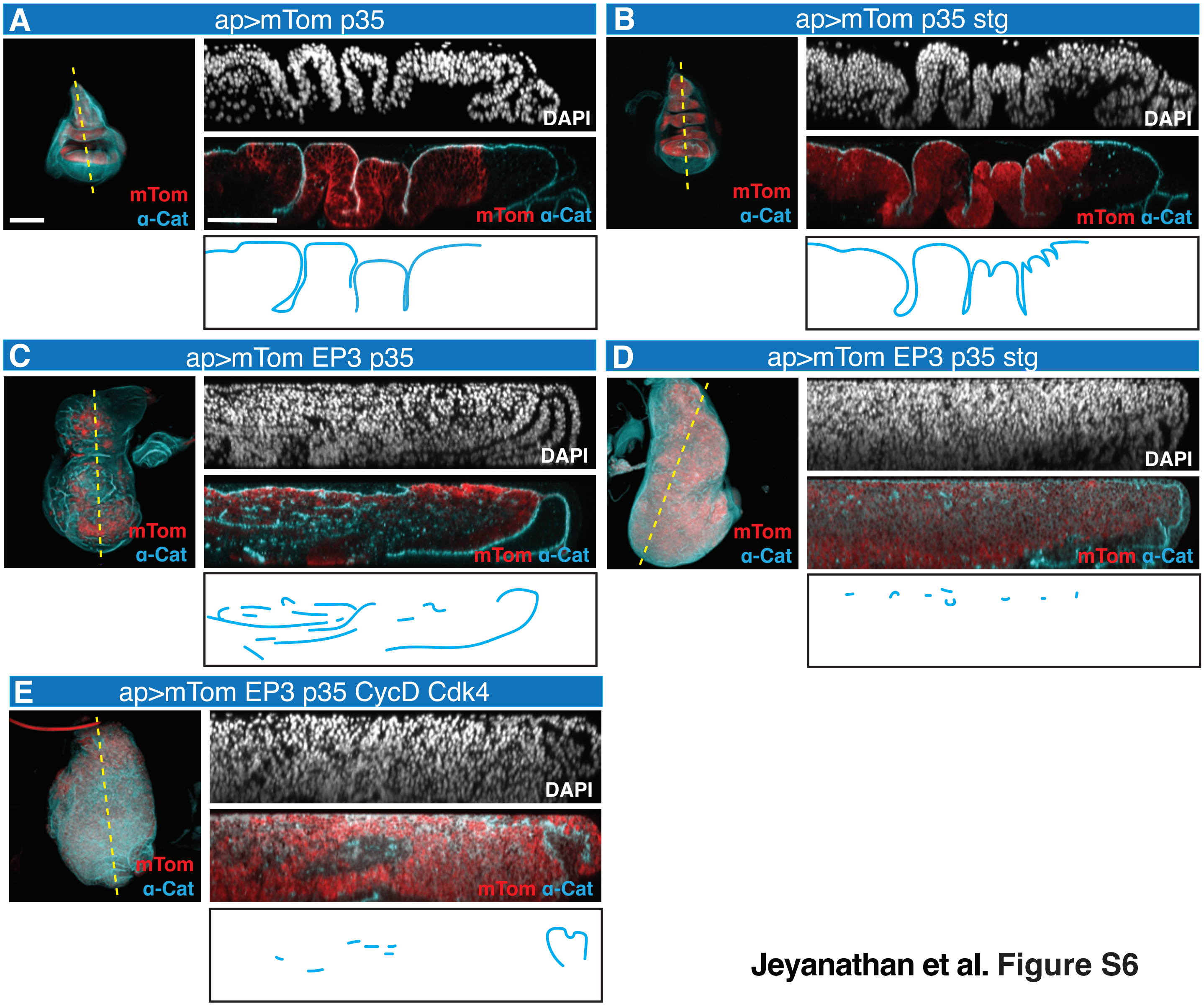

**Jeyanathan et al. Figure S6**

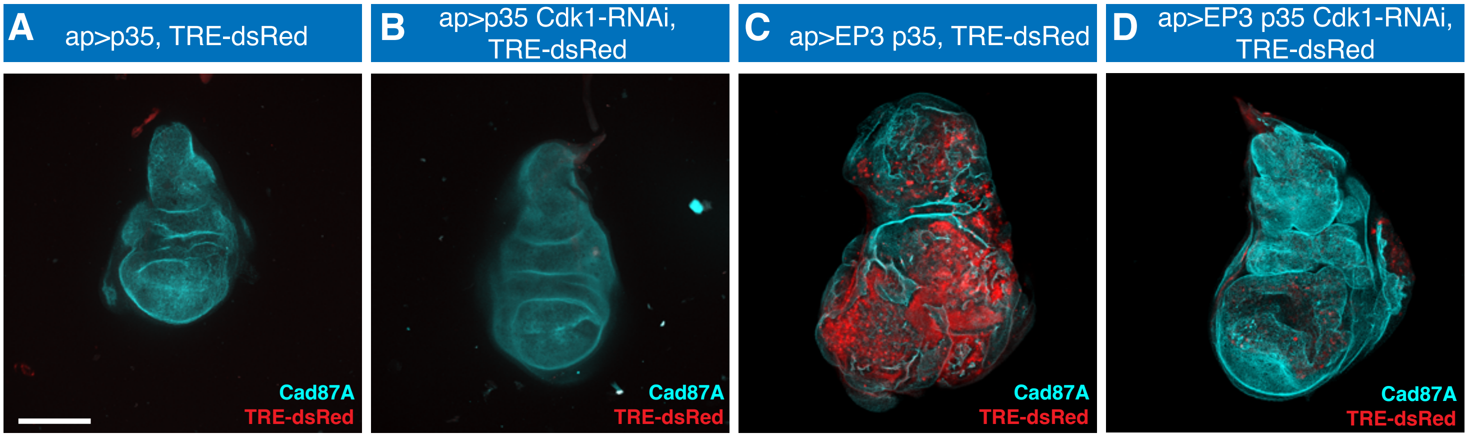

**Jeyanathan et al. Figure S7**

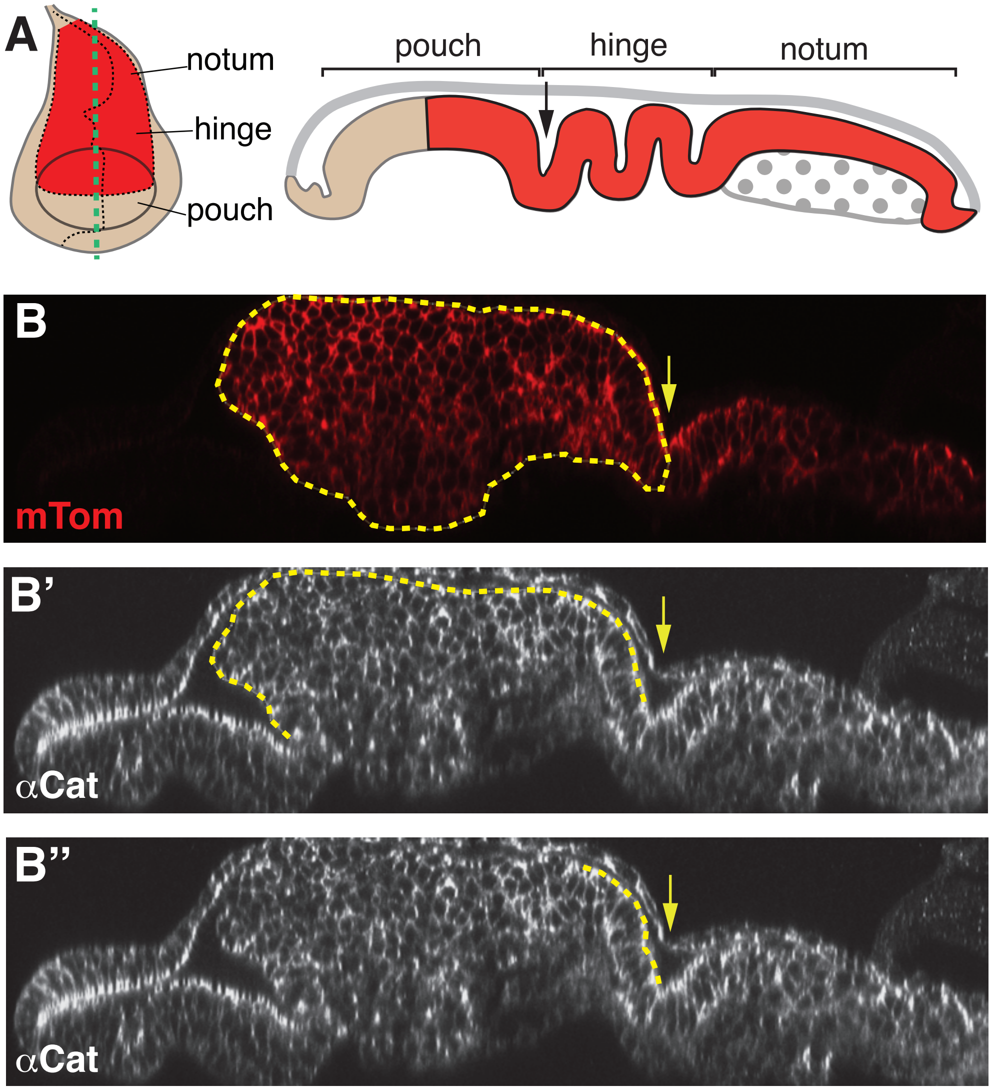

**Jeyanathan et al. Figure S8**

### Table S1: Fly lines, antibodies, and dyes

| Reagent or Resource | Source | Notes |
|---|---|---|
| **Fly Lines** | | |
| WT (OregonR) | BDSC # 5 | |
| *fzy*$^{1}$*, P{matα4-GAL-VP16}67/ CyO* | this work | |
| *da-Gal4, crb*$^{11A22}$*/ TM6B* | this work | |
| *da-Gal4, stg*$^{4}$*/ TM6B* | this work | |
| *crb*$^{11A22}$*, stg*$^{4}$*/ TM3, Ser* | this work | |
| *da-Gal4, crb, stg*$^{4}$*/ TM3, Ser* | this work | |
| *UAS-Cdc42-DN, stg*$^{4}$ | this work | |
| *UAS-Cdc42-DN;UAS-stg* | this work | |
| *shg*$^{R69}$*, UAS-stg/CyO* | this work | |
| *P{matα4-GAL-VP16}67/ CyO* | [72] | |
| *fzy*$^{1}$*/CyO* | BDSC # 2492 | |
| *His2Av-mRFP (II)* | BDSC # 23651 | |
| *endo-Crb::GFP* | [19] | |
| *endo-Crb::GFP, Gap43::mCherry* | [73] | |
| *Endo-Ecad::GFP* | [19] | |
| endo-Sdt::GFP (‘Std3GFP’ 363) | [74] | |
| *UAS-Notch*$^{intra}$*/ TM3, Sb* | Gift from Gary Struhl | |
| *UAS-Notch*$^{intra}$*; Sp/CyO* | Gift from Gary Struhl | |
| *crb*$^{11A22}$*/TM6B* | | |
| *stg*$^{4}$*/ TM3, Ser* | BDSC # 2500 | |
| *da-Gal4* | BDSC # 55850 | |
| *UAS-sqh, Gap43::mCherry (attp40)* | Gift from Adam Martin | |
| *Kr*$^{1}$*/CyO* | BDSC # 3494 | |
| *UAS-stg* | BDSC # 4777 | |
| *UAS-Cdc42DN* | BDSC # 6288 | |
| *shg*$^{R69}$*/CyO* | | |
| *ap-Gal4, TRE-DsRED/CyO; MKRS/TM6B, tub-Gal80*$^{ts}$ | this work | |
| *ap-Gal4, TRE-DsRED/CyO; UAS-p35/TM6B, tub-Gal80*$^{ts}$ | this work | |
| *ap-Gal4, UAS-Tomato/CyO; MKRS/TM6B, tub-Gal80*$^{ts}$ | this work | |

| | | |
|---|---|---|
| *en-Gal4, UAS-RFP (II) /CyO;MKRS/TM6B* | this work | |
| *UAS-E2F-PIP-3A, UAS-p35/TM6B, tub-Gal80$^{ts}$* | this work | |
| *UAS-scrib-RNAi, UAS-p35/TM6B, tub-Gal80$^{ts}$* | this work | |
| *ap-Gal4, UAS-myrTomato/CyO* | Gift from Marco Milan<br>BDSC # 32222 | |
| *en-Gal4,UAS-RFP/CyO* | BDSC # 30557 | |
| *sna/CyO;tub-Gal80$^{ts}$* | BDSC # 7018 | |
| *tub-Gal80$^{ts}$; TM2/TM6B* | BDSC # 7019 | |
| *UAS-Cdk1 RNAi (III)* | BDSC # 35350 | |
| *UAS-Cdk1 RNAi (III)* | BDSC # 28368 | |
| *UAS-Cdk1 RNAi (III)* | BDSC # 40950 | |
| *UAS-crb-RNAi (II)* | BDSC # 38373 | |
| UAS-*crb*-RNAi (III) | BDSC # 27697 | |
| *TRE-DsRED (II)* | BDSC # 59012 | |
| *UAS-scrib-RNAi (III)* | BDSC # 35748 | |
| *UAS-CycD,UAS-Cdk4/CyO* | Datar et al., 2019 | |
| *UAS-E2f1-PIP-3A (=EP3)* | [47] | |
| *UAS-LacZ (III)* | | |
| *UAS-p35 (III)* | BDSC # 5073 | |
| *UAS-stg (III)* | BDSC # 4778 | |
| **Primary antibodies** | | |
| anti-Yrt (guinea pig) | GP7; [75] | 1:500 |
| anti-Arm (mouse) | N2-7A1; DSHB | 1:50 |
| anti-Baz (rabbit) | Gift from Tony Harris | 1:3500 |
| anti-PH3 (rabbit) | 06-570; Millipore | 1:1000 |
| anti-PH3 (rat) | H9908; Sigma | 1:500 |
| anti-Sdt (rabbit) | [76] | 1:3500 |
| anti-DE-cad (rat) | DCAD2; DSHB | 1:20 |
| anti-Dlg (mouse) | 4F3; DSHB | 1:100 |
| Anti-Na/K-ATPase (mouse) | Nrv5F7; DSHB | 1:20 |
| anti-Cad87A (guinea pig) | Gift from D. Godt; [77] | 1:3000 (wing discs);<br>1:6000 |
| anti-Crb (rat) | F3; [46] | 1:1000 |
| anti-Dlg (mouse) | 4F3; DSHB | 1:20 |
| anti-Hemes (mouse) | [78] | |

| | | |
|---|---|---|
| anti-LanB1 (rabbit) | EPR3189; Abcam | 1:500 |
| anti-Mmp1 (mouse) | 3A6B4; 3B8D12; 5H7B11; DSHB | 1:20 |
| anti-Wg (mouse) | 4D4; DSHB | 1:50 |
| anti-α-Cat (guinea pig) | gp121; [79] | 1:500; 1:750 |
| **Alexa Fluor Secondary Antibodies** | | |
| Guinea pig, 488 | A11073; Thermo Fisher Scientific | 1:400 |
| Guinea Pig, 555 | A21435; Thermo Fisher Scientific | 1:400 |
| Guinea Pig, 647 | A21450; Thermo Fisher Scientific | 1:400 |
| Mouse, 488 | A11029; Thermo Fisher Scientific | 1:400 |
| Mouse, 555 | A21424; Thermo Fisher Scientific | 1:400 |
| Mouse, 647 | A21236; Thermo Fisher Scientific | 1:400 |
| Rat, 488 | A11006; Thermo Fisher Scientific | 1:400 |
| Rat, 555 | A10522/A21434; Thermo Fisher Scientific | 1:400 |
| Rat, 647 | A21247; Thermo Fisher Scientific | 1:400 |
| Rabbit, 488 | A11034; Thermo Fisher Scientific | 1:400 |
| Rabbit 555 | A21429; Thermo Fisher Scientific | 1:400 |
| Rabbit, 647 | A21245; Thermo Fisher Scientific | 1:400 |
| **Dyes** | | |
| DAPI | D3571Thermo Fisher Scientific | 1:1000 |
| Phalloidin-488 | PHDG1-A Cytoskeleton Inc. | 1:100 |
| CM-H2DCFDA | C6827; Thermo Fisher Scientific | 1:1000 |

BDSC: Bloomington Drosophila Stock Center
DSHB: Developmental Studies Hybridoma Bank

### Table S2: Genotypes and n-values

| **Embryo analysis** | | | |
|---|---|---|---|
| **Figure** | | **Genotype of embryo/tissue shown** | **n-value** |
| **Figure 1** | | | |
| **Figure 1A** | | wt | 10 |
| **Figure 1B** | | *fzy*$^{1}$/*fzy*$^{1}$ | 10 |
| **Figure 1C, D** | | *endo-Crb::GFP; His2Av-mRFP* | 3 embryos; 6-8 cells/embryo |
| **Figure 1E** | | *endo-Crb::GFP, Gap43::mCherry* | 3 embryos; 5 cells/embryo |
| **Figure 1E** | | *endo-Ecad::GFP* | 3 embryos; 4-7 cells/embryo |
| **Figure 1F, G** | | *endo-Ecad::GFP; crb* $^{11A22}$/ *crb* $^{11A22}$ | 3 embryos; 5-7 cells/embryo |
| **Figure 2** | | | |
| **Figure 2B-E** | wt | wt; *endo-Ecad::GFP* | 14 |
| | *crb* | *UAS-Nintra; endo-Ecad::GFP; crb*$^{11A22}$*, stg*$^{4}$*/ crb*$^{11A22}$ | 16 |
| | *stg crb* | *UAS-Nintra; endo-Ecad::GFP; crb*$^{11A22}$*, stg*$^{4}$*/ crb*$^{11A22}$*, stg*$^{4}$ | 4 |
| | *Ni stg crb* | *UAS-Nintra; endo-Ecad::GFP; crb*$^{11A22}$*, stg*$^{4}$*/ crb*$^{11A22}$*, stg*$^{4}$*, daGal4* | 11 |
| **Figure 2F** | wt | wt | 45 |
| | *crb* | *UAS-Nintra; crb*$^{11A22}$*, stg*$^{4}$*/ crb*$^{11A22}$ | 97 |
| | *stg* | *UAS-Nintra; crb*$^{11A22}$*, stg*$^{4}$*/ stg*$^{4}$ | 68 |
| | *Ni* | *UAS-Nintra; crb*$^{11A22}$*, stg*$^{4}$*/ daGal4* | 114 |
| | *stg crb* | *UAS-Nintra; crb*$^{11A22}$*, stg*$^{4}$*/ crb*$^{11A22}$*, stg*$^{4}$ | 74 |
| | *Ni crb* | *UAS-Nintra; crb*$^{11A22}$*, stg*$^{4}$*/ crb*$^{11A22}$*, daGal4* | 203 |
| | *Ni stg* | *UAS-Nintra; crb*$^{11A22}$*, stg*$^{4}$*/ stg*$^{4}$*, daGal4* | 125 |
| | *Ni stg crb* | *UAS-Nintra; crb*$^{11A22}$*, stg*$^{4}$*/ crb*$^{11A22}$*, stg*$^{4}$*, daGal4* | 171 |
| **Figure 3** | | | |
| **Figure 3A/3F** | wt | wt | 7/7 |
| **Figure 3B/3D** | *Cdc42-DN* | *UAS-Cdc42-DN; daGal4* | 10/3 |
| **Figure 3C** | *Cdc42-DN, stg* | *UAS-Cdc42-DN; stg*$^{4}$*/ stg*$^{4}$*, daGal4* | 8 |
| **Figure 3E** | *Cdc42-DN, Stg* | *UAS-Cdc42-DN; UAS-stg /daGal4* | 2 |
| **Figure 3G** | *shg* | *shg*$^{R69}$*/ shg*$^{R69}$ | 8 |
| **Figure 3H** | *shg, stg* | *shg*$^{R69}$*/ shg*$^{R69}$*; stg*$^{4}$*/ stg*$^{4}$ | 6 |
| **Figure 3I** | wt | wt | 34 |
| | *Cdc42-DN* | *UAS-Cdc42-DN; daGal4/ +* | 42 |
| | *Stg* | *UAS-stg /daGal4* | 75 |
| | *Cdc42-DN, stg* | *UAS-Cdc42-DN; stg*$^{4}$*/ stg*$^{4}$*, daGal4* | 71 |
| | *Cdc42-DN, Stg* | *UAS-Cdc42-DN; UAS-stg /daGal4* | 45 |

| | | | |
|---|---|---|---|
| **Figure S1** | | | |
| | $fzy^1/fzy^1$ | Sdt/Cad87A | 10 |
| | $fzy^1/fzy^1$ | Ecad/Dlg | 14 |
| | $fzy^1/fzy^1$ | aPKC::GFP/NaKAtpase | 3 |
| | $fzy^1/fzy^1$ | Par6::GFP/NaKAtpase | 3 |
| | $fzy^1/fzy^1$ | Yrt/Baz | 7 |
| **Figure S2** | | | |
| | | endo-Sdt::GFP (Std3GFP 363) | 6 embryos |
| **Figure S3** | | | |
| | wt | wt | 45 |
| | *crb* | *UAS-Nintra;* $crb^{11A22}$, $stg^4$/ $crb^{11A22}$ | 97 |
| | *Ni* | *UAS-Nintra;* $crb^{11A22}$, $stg^4$/ *daGal4* | 114 |
| | *stg* | *UAS-Nintra;* $crb^{11A22}$, $stg^4$/ $stg^4$ | 68 |
| | *Ni crb* | *UAS-Nintra;* $crb^{11A22}$, $stg^4$/ $crb^{11A22}$, *daGal4* | 203 |
| | *Ni stg* | *UAS-Nintra;* $crb^{11A22}$, $stg^4$/ $stg^4$, *daGal4* | 125 |
| | *Ni stg crb* | *UAS-Nintra;* $crb^{11A22}$, $stg^4$/ $crb^{11A22}$ , $stg^4$, *daGal4* | 171 |
| | *Kr* | *Kr/Kr* | 20 |
| | *Kr stg crb* | *Kr/Kr;* $stg^4$, $crb^{11A22}$/ $stg^4$, $crb^{11A22}$ | 31 |
| | *Cdc42-DN* | *UAS-Cdc42-DN; daGal4* | 42 |
| | *Cdc42-DN Stg* | *UAS-Cdc42-DN, UAS-stg, daGal4* | 45 |
| | *BazSA::GFP* | *UAS-BazSA::GFP/ daGal4* | 47 |
| | *BazSA::GFP Ni stg* | *UAS-BazSA::GFP, UAS-Nintra,* $stg^4$/ $stg^4$, *daGal4* | 83 |
| **Figure S4** | | | |
| | wt | wt | 10 |
| | *Kr* | *Kr/Kr* | 4 |
| | *crb* | $crb^{11A22}$/ $crb^{11A22}$ | 8 |
| | *Ni Kr crb* | *UAS-Nintra ; Kr/ Kr;* $crb^{11A22}$, $stg^4$/ $crb^{11A22}$, *daGal4* | 13 |
| | *Kr stg crb* | *UAS-Nintra; Kr/ Kr;* $crb^{11A22}$, $stg^4$/ $crb^{11A22}$, $stg^4$ | 6 |

| **Imaginal wing disc analysis** | | | |
|---|---|---|---|
| **Figure** | **Genotype** | **n value** | **Notes** |
| **Figure 4** | | | |
| **Figure 4B** | *w; ap-Gal4, UAS-myrTomato; UAS-p35/UAS-lacZ* | 12 | Larvae raised at 18ºC |
| | *w; en-Gal4, UAS-RFP; UAS-crb-RNAi* | 30 | Larvae raised at 25ºC |
| **Figure 4C** | *w; ap-Gal4, UAS-myrTomato; UAS-E2F1::PIP3, UAS-p35/UAS-lacZ* | 30 | Larvae raised at 18ºC |
| **Figure 4D** | *w; ap-Gal4, UAS-myrTomato; UAS-E2F1::PIP3, UAS-p35/UAS-crb-RNAi* | 40 | Larvae raised at 18ºC |
| **Figure 4E** | *w; en-Gal4, UAS-RFP/+; UAS-p35/+* | 10 | Larvae raised at 29ºC |
| | *w; en-Gal4, UAS-myrTomato; UAS-E2F1::PIP3, UAS-p35* | 15 | Larvae raised at 29ºC |
| | *w;* en-*Gal4, UAS-myrTomato; UAS-E2F1::PIP3, UAS-p35/ UAS-crb-RNAi* | 15 | Larvae raised at 29ºC |

| | | | |
|---|---|---|---|
| **Figure 4F** | *w; ap-Gal4, UAS-myrTomato; UAS-p35/ UAS-lacZ* (control) | 12 | Larvae raised at 29ºC |
| | *w; ap-Gal4, UAS-myrTomato; UAS-E2F1::PIP3, UAS-p35/UAS-lacZ* | 12 | Larvae raised at 29ºC |
| | *w; ap-Gal4, UAS-myrTomato; UAS-E2F1::PIP3, UAS-p35/UAS-crb-RNAi* | 12 | Larvae raised at 29ºC |
| **Figure 4G** | *w; ap-Gal4, UAS-myrTomato; UAS-crb-RNAi* (control) | 7 | Larvae raised at 18ºC or 29ºC and dissected at 3 days old 3rd instar |
| | *w; ap-Gal4, UAS-myrTomato; UAS-E2F1::PIP3, UAS-p35/UAS-lacZ* | 15 | Larvae raised at 18ºC and dissected at 3 days old 3rd instar |
| | *w; ap-Gal4, UAS-myrTomato; UAS-E2F1::PIP3, UAS-p35/UAS-lacZ* | 10 | Larvae raised at 18ºC and dissected at 5 days old 3rd instar |
| | *w; ap-Gal4, UAS-myrTomato; UAS-E2F1::PIP3, UAS-p35/UAS-lacZ* | 10 | Larvae raised at 29ºC and dissected at 3 days old 3rd instar |
| | *w; ap-Gal4, UAS-myrTomato; UAS-E2F1::PIP3, UAS-p35/UAS-crb-RNAi* | 12 | Larvae raised at 18ºC and dissected at 3 days old 3rd instar |
| **Figure 4H** | *w; ap-Gal4, UAS-myr Tomato; UAS-p35/ UAS-lacZ* (control) | 62 | Larvae raised at 18ºC |
| | *w; ap-Gal4, UAS-myrTomato; UAS-E2F1::PIP3, UAS-p35/UAS-lacZ* | 263 | Larvae raised at 18ºC |
| | *w; ap-Gal4, UAS-myrTomato; UAS-E2F1::PIP3, UAS-p35/UAS-crb-RNAi* | 261 | Larvae raised at 18ºC |
| **Figure 5** | | | |
| **Figure 5A** | *w; ap-Gal4, UAS-myrTomato/tub-Gal80ts; UAS-p35/ UAS-lacZ* | 10 | Larvae raised at 18ºC and moved to 29 ºC at third instar to inactivate Gal80ts for 72 hours |
| **Figure 5B** | *w; ap-Gal4, UAS-myrTomato/tub-Gal80ts; UAS-p35/ UAS-Cdk1-RNAi* | 14 | Larvae raised at 18ºC and moved to 29ºC at third instar to inactivate Gal80ts |
| **Figure 5C** | *w; ap-Gal4, UAS-myrTomato/tub-Gal80ts; UAS-scrib-RNAi, UAS-p35/ UAS-lacZ* | 16 | Larvae raised at 18ºC and moved to 29ºC at 3rd instar to inactivate Gal80ts |
| **Figure 5D** | *w; ap-Gal4, UAS-myrTomato/tub-Gal80ts; UAS-scrib-RNAi, UAS-p35/ UAS- Cdk1-RNAi* | 15 | Larvae raised at 18ºC and moved to 29ºC at third instar to inactivate Gal80ts |
| **Figure S5** | | | |
| | *w; ap-Gal4, UAS-myrTomato; UAS-p35* | shown in the figure | Larvae raised at 25ºC or 29ºC and dissected when >50% have pupariated |
| | *w; ap-Gal4, UAS-myrTomato; UAS-E2F1::PIP3, UAS-p35* | shown in the figure | Larvae raised at 25ºC or 29 ºC |
| **Figure S6** | | | |
| **Figure S6A** | *w; ap-Gal4, UAS-myrTomato; UAS-p35/UAS-lacZ* | 25 | Larvae raised at 29ºC and dissected when >50% have pupariated |
| **Figure S6B** | *w; ap-Gal4, UAS-myrTomato/ UAS-stg; UAS-p35* | 15 | Larvae raised at 29ºC |
| **Figure S6C** | *w; ap-Gal4, UAS-myrTomato; UAS-E2F1::PIP3, UAS-p35/UAS-lacZ* | 30 | Larvae raised at 29ºC |
| **Figure S6D** | *w; ap-Gal4, UAS-myrTomato/ UAS-stg; UAS-E2F1::PIP3, UAS-p35* | 10 | Larvae raised at 29ºC |
| **Figure S6E** | *w; ap-Gal4, UAS-myrTomato/ UAS-CycD UAS-Cdk4; UAS-E2F1::PIP3, UAS-p35* | 12 | Larvae raised at 29ºC |

| Figure S7 | | | |
|---|---|---|---|
| **Figure S7A** | *w; ap-Gal4, TRE-dsRed; UAS-p35/UAS-lacZ* | 15 | Larvae raised at 25ºC and dissected once >70% have pupariated |
| **Figure S7B** | *w; ap-Gal4, TRE-dsRed; UAS-p35/UAS-Cdk1-RNAi* | 10 | Larvae raised at 29ºC |
| **Figure S7C** | *w; ap-Gal4, TRE-dsRed ; UAS- E2F1::PIP3, UAS-p35/UAS-lacZ* | 30 | Larvae raised at 29ºC |
| **Figure S7D** | *w; ap-Gal4, TRE-dsRed; UAS- E2F1::PIP3, UAS-p35/UAS-Cdk1-RNAi* | 11 | Larvae raised at 29ºC |
